# MMAD-Risk: Multivariate Mixed Survival Analysis for the Prediction of Age-Dependent Disease Risks from Plasma Proteomes

**DOI:** 10.64898/2026.09.16.752074

**Authors:** Amelie M. Hilger, Johannes Söding

## Abstract

**Motivation:** Multivariate survival analysis with hundreds of correlated outcomes is computationally challenging. Established approaches either ignore correlations between response variables, rely on black-box deep learning or are limited to small-scale outcomes.

**Results:** We introduce MMAD-Risk, a novel multivariate mixed accelerated failure time (AFT) model that enables scalable analysis of high-dimensional survival analysis data. We train MMAD-Risk using amortized variational inference where we design the variational distribution such that it factorizes across diseases, allowing us to decompose multivariate disease risk prediction into a series of tractable, one-dimensional problems. This allows us to calculate the ELBO analytically, enabling fast computation. The model employs a low-rank decomposition of the effect size matrix **B** = **VW**^⊤^ to capture shared disease mechanisms and latent random effects **Vz**_*n*_ to model comorbidity. MMAD-Risk is trained on the UK Biobank Pharma Proteomics cohort (*N* ≈ 55000, *P* ≈ 3000 proteins, *D* = 271 diseases). Using the full 3,000-protein dataset, MMAD-Risk achieved a mean concordance index (c-index) of 0.744 for diagnoses occurring ⩾ 10 years after blood sample collection, outperforming a Cox proportional hazards model (mean c-index = 0.709). Greedy backward selection identified a 10-protein panel that preserved *>* 99% of the full-model performance. On this reduced panel MMAD-Risk still outperformed Cox regression (0.738 vs. 0.679).

**Availability and implementation:** https://github.com/soedinglab/MMAD-Risk.git

**Supplementary information:** Supplementary data are available at *Bioinformatics* online.

## Introduction

Ageing is the leading risk factor for various different diseases, such as cardiovascular disease, neurodegenerative diseases or cancer (Niccoli and Partridge, 2012; Tenchov et al., 2023; Partridge et al., 2018). As the word population is ageing (Beard et al., 2016), age-dependent diseases contribute increasingly to the overall global disease burden. In order to address this health crisis, we need good models of ageing. One model of ageing are ageing clocks (Rutledge et al., 2022; Argentieri et al., 2024; Horvath, 2013; Hannum et al., 2013; Lehallier et al., 2019; Sedaghat et al., 2025), which train a machine learning model on biological data such as epigenetic or proteomic data to predict the study participants’ age. This predicted age is then taken as a proxy for a biological age. Based on the difference between chronological age and biological age an “age gap” is calculated, where a higher biological than chronological age is associated with a higher risk of getting diagnosed with various different diseases. Even though ageing clocks are a helpful tool to understand ageing and age-related disease, they collapse ageing into a single number that does not account for different systems ageing at different rates.

Organ ageing clocks address this shortcoming (Oh et al., 2023, 2025; Tian et al., 2023; Li et al., 2025b; MULTI Consortium et al., 2025; Wang et al., 2025a; Yu et al., 2025; Wang et al., 2025b; Wen, 2025; Kivimäki et al., 2025; Goeminne et al., 2025). They are trained on data specific to individual organs, for example organ-specific imaging data or only on plasma proteins that are much more strongly expressed in one organ than in any other organ. These clocks predict organ specific disease risk more accurately than an ageing clock that predict only a single biological age. However, organ ageing clocks do not consider shared disease mechanisms across different organ systems.

In 2023, UK Biobank published the largest plasma proteomics data set available so far, containing plasma protein concentrations for almost 3000 proteins and 55000 study participants (Sun et al., 2023). This data set has enabled rapid progress in various areas, such as the creation of a new human proteome atlas (Deng et al., 2025), multi-disease models (Carrasco-Zanini et al., 2024; Gadd et al., 2024; You et al., 2023), and a better understanding of cardiovascular and neurodegenerative diseases (Li et al., 2025a; Guo et al., 2024).

In contrast to only looking at single diseases, multimorbidity models take several diseases into account at once: Shmatko et al. (2025) have developed a large health model that is able to predict future health events similar to a large language model, Jiang et al. (2025) and You et al. (2023) have developed multivariate models where shared disease mechanisms are modeled by a neural network. Multivariate survival analysis models are suitable for the analysis of multimorbidity. These methods have been developed based on deep learning (Jiang et al., 2025; Tjandra et al., 2021; Lee et al., 2018) or graphs (Lillelund et al., 2026), while other approaches restrict the effect size matrix, for example by restricting its rank (Wang et al., 2017) or placing a sparse group lasso prior on it (Li et al., 2021). Maia et al. took a more classical approach by developing a multivariate mixed Cox proportional hazard model (Maia et al., 2014).

Hui et al. (2017) introduced a framework for Gaussian linear latent variable models (GLLVMs) suitable for Bernoulli, mutinomial and Poisson response variables. They factorized the multivariate response function into a product of one-dimensional response functions that are coupled via a latent multivariate normal distribution and employ a variational approximation with a multivariate normal distribution as variational distribution.

In this study, we present MMAD-Risk (Multivariate Mixed survival analysis for Age-dependent Disease risk) to predict age-dependent disease risks from plasma proteomes on UK Biobank. We model comorbidities by introducing a latent variable that models shared, latent effects across diseases that are unaccounted for in the proteome data, mirroring the model structure proposed by (Hui et al., 2017) and adapting it to survival analysis. Shared disease mechanisms are captured by a low-rank decomposition of the effect size matrix that models the influence of proteins on disease risk.

We train MMAD-Risk using amortized variational inference and select the variational distribution in a way that allows us to decompose the complex, high dimensional disease risk prediction into a series of one-dimensional problems. We can calculate the evidence lower bound (ELBO) of MMAD-Risk analytically, enabling the fast computation of disease risk.

MMAD-Risk is the first instance of a multivariate mixed survival analysis model being applied to over 100 outcomes, and the first to be applied to the UK Biobank Plasma Proteomics Data Set. In contrast to ageing clocks, MMAD-Risk does not rely on the difference between predicted biological and chronological age. It is able to model how the same biological processes influence disease risk across different organ systems. As a generalized linear model, its effect sizes are directly interpretable, allowing us to identify the most relevant proteins for disease risk prediction.

## Methods

### MMAD-Risk

Suppose we have *N* patients in a population-based study such as UK Biobank. The study records the diagnoses of patients for *D* different diseases. Let *δ*_*nd*_ ∈ {0, 1} be the censoring variable indicating whether patient *n* has already been diagnosed with disease *d* ∈ {1, …, *D*} and *y*_*nd*_ the age at diagnosis, or the current age if the disease has not been diagnosed yet (*δ*_*nd*_ = 0). We also have *P* covariates **x**_*n*_ ∈ ℝ^*P*^ for each patient such as current age, sex, BMI etc., and information derived from proteomic measurements of thousands of blood protein concentrations (Sun et al., 2023; Argentieri et al., 2024).

We estimate the biological age of a patient by multiplying the chronological age *y* with a patient- and disease-specific acceleration/deceleration factor 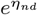. Here, *η*_*nd*_ can be estimated as fixed effect from the covariates, 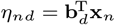, with regression weights **b**_*d*_. The log likelihood for patient *n* in a univariate accelerated failure time (AFT) model is given by

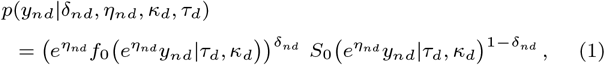

where *S*_0_(*t*|*κ, τ*) = exp − (*t/τ*)^*κ*^ is the Weibull survival function representing the patients who did not yet get disease *d* before their current chronological age *y*_*nd*_, and patients who got disease *d* at age *y*_*nd*_ are represented by the corresponding Weibull density 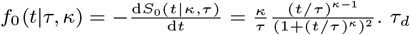 and *κ* are the Weibull scale and shape parameters with respect to diseases *d*.

Here, we introduce a multivariate accelerated failure time model, in which, in addition to the *P* fixed effects **Bx**_*n*_, *K* random effects **z** ~ *N*(**z**|**0, *I***_*K*_) couple the failure rates of the *D* diseases, mirroring the approach introduced by Hui et al. (2017):

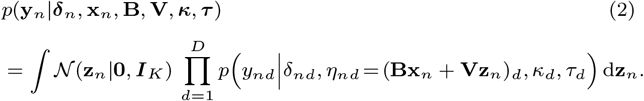

The fixed-effects matrix **B** = **VW**^*⊤*^ contains the regression weights for the covariates **X** = (**x**_1_, …, **x**_*N*_), where **B** ∈ ℝ^*D×P*^, **V** ∈ ℝ^*D×K*^ and **W** ∈ ℝ^*P ×K*^. The *K* latent dimensions represent *factors of ageing*, which represent different ageing mechanisms that are shared across diseases. The matrix **W** maps the contribution of plasma protein concentration to the factors of ageing, whereas **V** maps the influence of the factors of ageing on the diseases. The latent variables **z**_*n*_ model comorbidities, correlations in risk between diseases due to unknown, latent factors that are not accounted for by the covariates **X**. This approach of treating the survival times with respect to different diseases as independent and coupling them via a multivariate normal distribution is similar to the approach in (Callegher et al., 2024). We use amortized variational inference to train MMAD-Risk.

### Variational Inference

Variational inference is a class of techniques for approximating intractable probability distributions and the integrals associated with Bayesian inference. (Bishop, 2006). Given a posterior probability distribution *p*(**z**|**y, x, *θ***) over a latent variable **z** we aim to approximate this distribution with a variational distribution *q*(**z**|***ϕ***) depending on a variational parameters ***ϕ***. We aim to select a variational distribution *q*(**z**|***ϕ***) that can approximate *p*(**z**|**y, x, *θ***) well. The model parameters ***θ*** and variational parameters ***ϕ*** are optimized together by maximizing the evidence lower bound (ELBO):

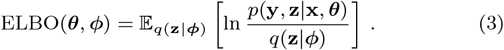

One can show that the ELBO is a lower bound to the log evidence ln *p*(**y**|**x, *θ***):

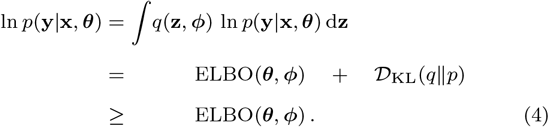

The Kullback Leibler divergence between two distribution is always non-negative and is a measure of similarity between the two distributions. By optimizing the ELBO we make the variational distribution as similar as possible to the posterior distribution and at the same time we optimize the lower bound of the evidence, thereby maximizing the evidence when the lower bound is tight.

### Training MMAD-Risk with variational inference

The likelihood over the multivariate survival analysis function is given by equation (2). Since this integral is intractable, we need to employ variational inference to approximate the likelihood. To obtain a good approximation with as few variational parameters as possible, we design the form of our variational distribution such that it mirrors the structure of the marginal likelihood in equation (2):

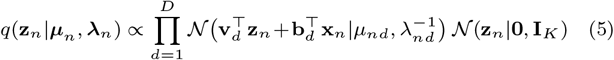

The normal distribution *N*(**z**_*n*_|**0, I**_*K*_) is shared between the marginal likelihood of the model and the variational distribution. Furthermore, we approximate the function over *η*_*nd*_, *p*(*y*_*nd*_ |*δ*_*n d*_, *η*_*nd*_, *κ*_*d*_, *τ*_*d*_), up to a constant factor, by a normal distribution. The variational parameters ***μ***_*n*_ = (*μ*_*n*1_, …, *μ*_*nD*_) and ***λ***_*n*_ = (*λ*_*n*1_, …, *λ*_*nD*_) are learned from the data. Crucially, by selecting the variational distribution in this way, we show in the Supplementary Material that the ELBO can be calculated analytically:

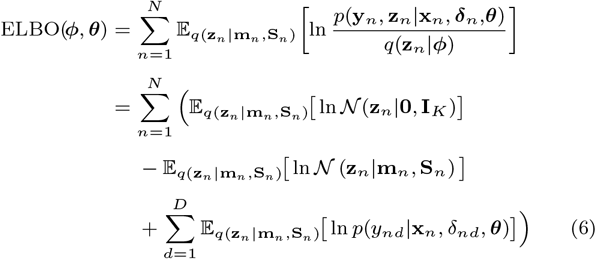

where

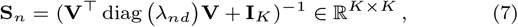

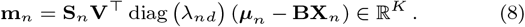

These three terms can be further transformed to

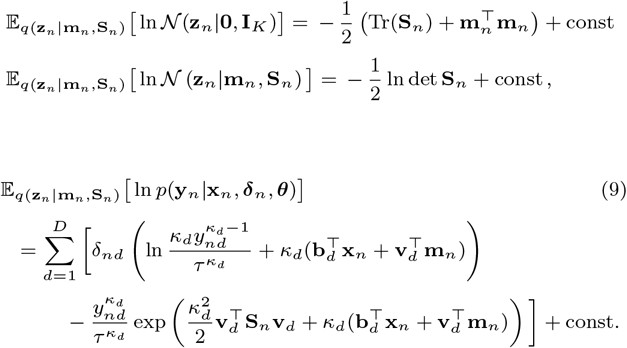

Since the ELBO and its gradient can be calculated analytically, it can be optimized very efficiently. We use ADAM implemented in Pytorch. One training run of MMAD-Risk with *K* = 15 and on all 3000 proteins takes 25 minutes for 200 epochs on a mem1 hdd1 x36 instance on the UK Biobank Research Analysis Platform. We set the learning rate to 0.001.

To avoid reverse-causation, we introduce a mask that removes all diagnoses that were given before blood sample collection. Information on this and on the regularization of MMAD-Risk is given in the Supplementary Information.

### Amortization

To prevent overfitting, we use amortization. Instead of fitting ***μ***_*n*_ and ***λ***_*n*_ directly as the variational parameters, we train a neural network to predict (*μ*_*nd*_, *λ*_*nd*_) from the age at onset *y*_*nd*_ and disease status *δ*_*nd*_,

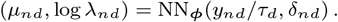

and learn its weights ***ϕ*** as variational parameters. We can see from eq. (1) that *p y*_*nd*_ *δ*_*nd*_, *η*_*nd*_, *κ*_*d*_, *τ*_*d*_ depends on *y*_*nd*_ only through *y*_*nd*_*/τ*_*d*_, which is why we use it as input to our neural network. For the analysis of the full panel with 3000 proteins, we use a neural network with two input and two output nodes and three hidden layers with 128 nodes each. For the prediction on the 10-protein panel, we use two hidden layers with 128 nodes each. In both cases, we use the hyperbolic tangent as activation function.

### Time complexity

The computation of the matrices 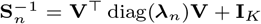 takes *O*(*NDK*^2^), and the inversion and calculation of their determinants both take *O*(*NK*^3^). For the calculation of **m**_*n*_, we need to compute 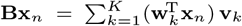 which takes in *O*(*NKP* + *NKD*). The calculation of **S**_*n*_**V**^*⊤*^ diag(***λ***_*n*_)**(*μ***_*n*_ − **Bx**_*n*_) takes *O*(*NDK*^2^), hence the computation of **m**_*n*_ takes *O*(*NDK*^2^ + *NKP*) in total. For the calculation of the third ELBO term in equation (9), we need to calculate **Bx**_*n*_ in *O*(*NKP* +*NKD*), **v**_*d*_**m**_*n*_ in *O*(*NDK*) and 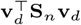 in *O*(*NDK*^2^). Since *K* is 15, *D* is 271, and *P* is ~ 3000, *P* dominates over *K*^2^ and *D*. Therefore, the overall complexity is *O*(*NKP*).

In the plasma proteomics data set in UK Biobank we have *N* ≈ 55 000 study participants and *P* = 2 923 plasma proteins, and *K* ≈ 15 factors of ageing, resulting in *NKP* ≈ 2.4 × 10^9^.

### Data Preprocessing

#### Calculating the protein gap

Protein concentrations may tend to increase or decrease with age (Álvez et al., 2025). For disease risk prediction, we want to capture the UK Biobank participants’ deviation from the plasma protein concentration that is typical for their age. We first fit a spline to the mean age-dependent protein concentration of UK Biobank participants, ***π***_*p*_ = *f*_*p*_(***a***) + ***ε***, *p* = 1, …, *P*, where ***a*** is the age at which the blood sample was collected and *f*_*p*_ is a spline with six nodes. We then calculate the residuals Δ***π***_*p*_ = ***π***_*p*_ − *f*_*g*_(***a***) and fit another spline to the average age-dependent variance of the plasma protein concentrations (Δ***π***_*p*_)^2^ = *g*_*p*_(***a***) + ***ε***, (*p* = 1, …, *P*), where *g*_*p*_ is another spline. Finally, we compute the normalized age-dependent protein gap as 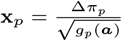 (Supplementary Figures 1 and 2).

#### Definition and inclusion of diseases

All 271 diseases included in the analysis were determined by taking diagnoses as a three digit ICD-10 code. Diagnoses with longer codes were merged with the corresponding three digit ICD-10 code. Diseases included in the analysis were selected from the first 15 chapters of the ICD-10 and had at least 150 incident cases within 10 years after the blood sample collection from UK Biobank participants. Figure 1A shows an overview of the diseases included in this study by their chapters in the ICD-10.

**Figure 1.**
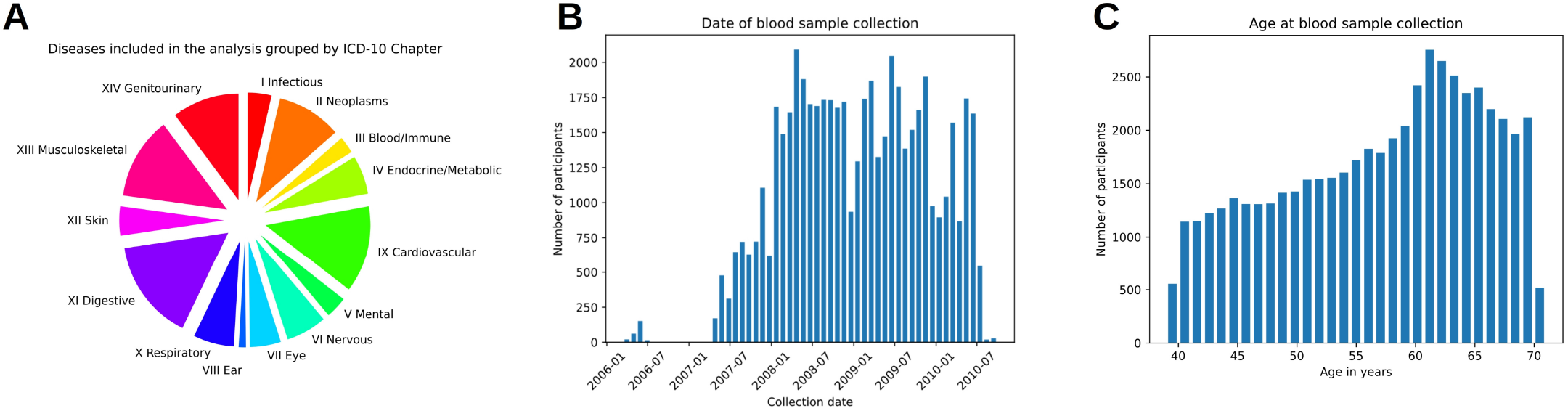
Information about the data included in the analysis. (A) Overview about the diseases included in the analysis, grouped by International Classification of Diseases 10th Revision (ICD-10) chapter. (B) Date of blood sample collecttion and (C) age in years on the day of blood sample collection for participants of the UK Biobank proteomics study.

#### Input Data

Besides the diseases and the protein gap data, we input BMI, sex and the age at blood sample extraction ***a*** into the model. Here, ***a*** gets transformed into a non-linear basis by inputting (*σ*(0.1(*a*_*n*_ − 40)), *σ*(0.1(*a*_*n*_ − 50)), *σ*(0.1(*a*_*n*_ − 60)), *σ*(0.1(*a*_*n*_ − 70))), where *σ*(*x*) = 1*/*(1 + exp(−*x*)) is the logistic sigmoid function.

## Results

### The UK Biobank pharma proteomics project

We train MMAD-Risk on the UK Biobank pharma proteomics project by Sun et al. (2023). We split the data set into independent training, validation and test data (*N*_train_ = 40000, *N*_val_ = 6496, *N*_test_ = 6496). The selection of all diseases with at least 150 incident cases within 10 years after blood sample collection results in a total of *D* = 271 being included into the analysis (Figure 1A). The date at which the blood samples for the plasma proteome measurements were collected from UK Biobank participants and their age at collection is shown in panels (B,C) of Figure 1. All the results reported in this section were derived before the shutdown of the UK Biobank Research Analysis Platform (UKB-RAP).

### Model Evaluation

We assign all study participants to the training, validation and test set without overlap between these sets. As evaluation method, we use the c-index, which measures the proportion of concordant pairs. A concordant pair is a pair of patients for which the person assigned with the higher disease risk got the disease earlier, while ignoring pairs for which both patients have not gotten the disease yet. (Li et al., 2022; Harrell et al., 1982):

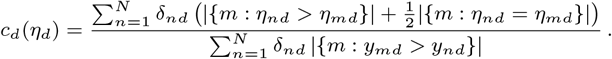

In all subsequent analyses, the c-index is evaluated on the diseases that were diagnosed ⩾ 10 years after the collection of the blood sample from which the plasma protein concentrations were measured. The models are trained on the training set using all diseases diagnosed after blood sample collection. They are validated and tested on all diseases diagnosed after ⩾ 10 years of the blood sample collection on the validation and test set, respectively (Figure 2A). We select *K* = 15 for MMAD-Risk and provide more details and in the supplement. Supplementary Figure 3 shows the performance of MMAD-Risk dependent on *K*.

### Model Comparison

We compare MMAD-Risk to a Cox proportional hazards model (Figure 2B) and a multivariate fixed model with ***η*** = **BX** = **VW**^*⊤*^**X** and design matrix **X** = (**x**_1_, …, **x**_*N*_). The multivariate fixed model uses the low-rank matrix decomposition of **B**, but it does not contain the mixed effect **Vz**_*n*_ and therefore does not take the covariance structure into account.

**Figure 2.**
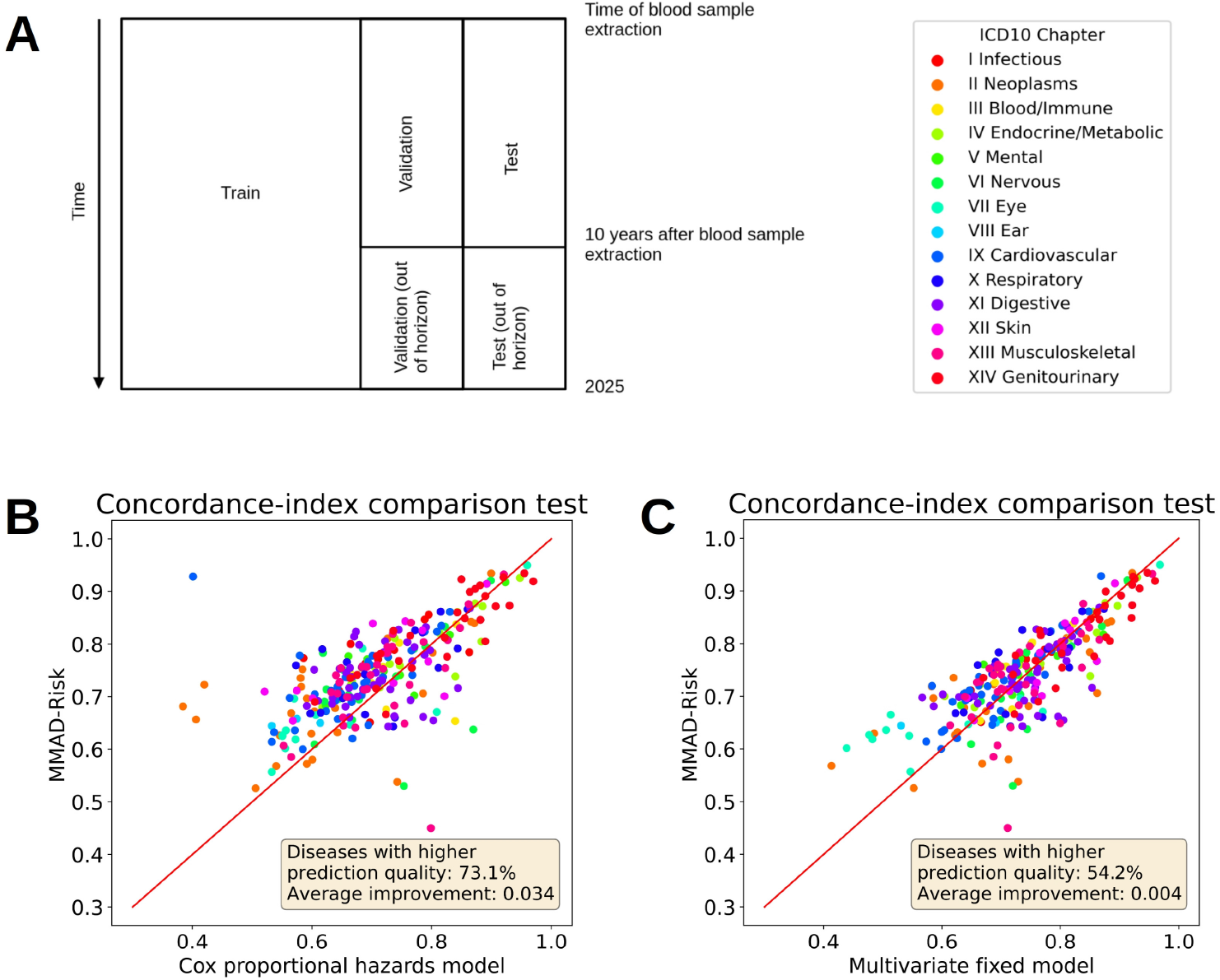
Model comparison on 3000 proteins. Model performance measured by c-index. (A) Scheme of evaluating the model performance during training, validating and testing. The data is split into disjoint training, validation and test set. The model is trained on all diseases diagnosed after blood sample collection in the training set. For validating and testing, the diagnoses between blood sample collection and 10 years after blood sample collection get are put into the model and the c-index is calculated on the predictions after these 10 years. Panels (B) and (C) display model performance comparison based on the c-index between MMAD-Risk and a Cox proportional hazards model and MMAD-Risk and a multivariate fixed survival analysis model, respectively.

MMAD-Risk outperforms the Cox proportional hazards model on 73.1% of diseases and improves the mean c-index over all 271 diseases by 0.034 (Figure 2B). In contrast, MMAD-Risk outperforms the multivariate fixed model only on 54.2% of diseases, achieving an improvement of the mean c-index by 0.004 (Figure 2C).

A comparison of MMAD-Risk with a univariate mixed effect model and logistic regression is shown in Supplementary Figure 4. Additionally, we compare the risk stratification of MMAD-Risk and the Cox model by plotting 1-Kaplan-Meier estimator for the top, medium and bottom risk decile (Supplementary Figure 10) (Kaplan and Meier, 1958).

### Detecting Disease Clusters

We cluster the entries of **V**, the matrix that maps the influence of the factors of ageing onto the 271 diseases in order to identify diseases with similar disease mechanisms. For this purpose, we apply hierarchical clustering using Ward’s method both to the diseases and to the factors of ageing. Based on the elbow method (Supplementary Figure 5), we identify 10 different clusters of diseases. The most important proteins for each of these clusters can be found in Figure 3B. Figure 5 (E) displays a comparison of which of the top 3 proteins of each cluster also appear in major proteomic studies (Argentieri et al., 2024; Deng et al., 2025; Carrasco-Zanini et al., 2024). The full dendrogram with disease labels is shown in Supplementary Figure 6. Further information on the normalization of **V** is given in the Supplementary Information as well.

**Figure 3.**
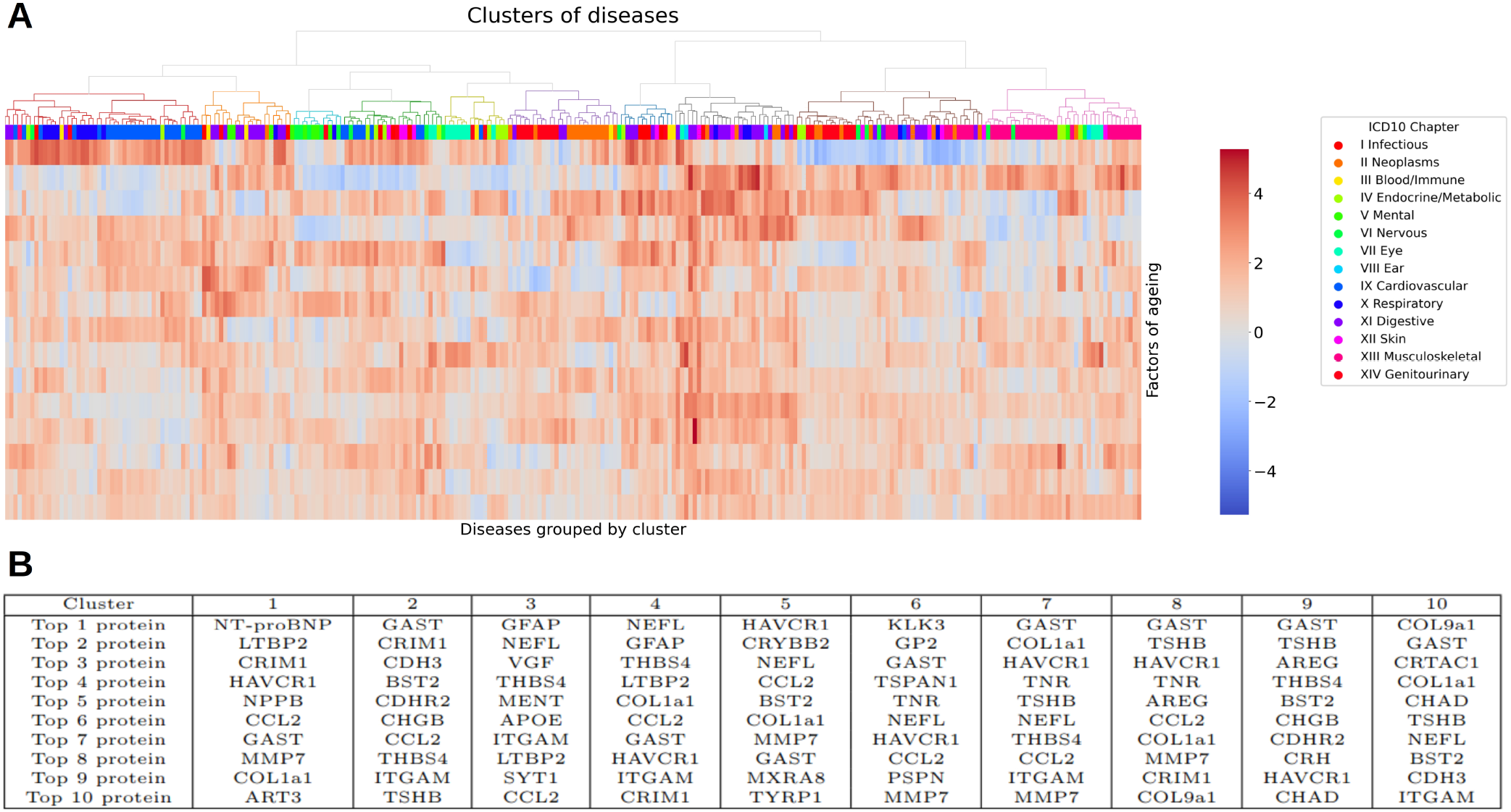
Clustering of diseases. (A) Heatmap of **V. V** is the matrix that maps the *K* factors of ageing onto disease risk. Factors of ageing represent distinct ageing processes that are shared across diseases. Diseases were clustered with hierarchical clustering using Ward’s method. Different clusters are color-coded on the dendrogram, diseases are color-coded by ICD-10 Chapter. (B) Ten most predictive protein in each cluster.

The covariance matrix of diseases **VV**^*⊤*^ is shown in Figure 4. Using agglomerative clustering with Ward’s method, we discover seven disease clusters. The elbow plot for these clusters is shown in Supplementary Figure 7, the full dendrogram with disease labels is shown in Supplementary Figure 78

**Figure 4.**
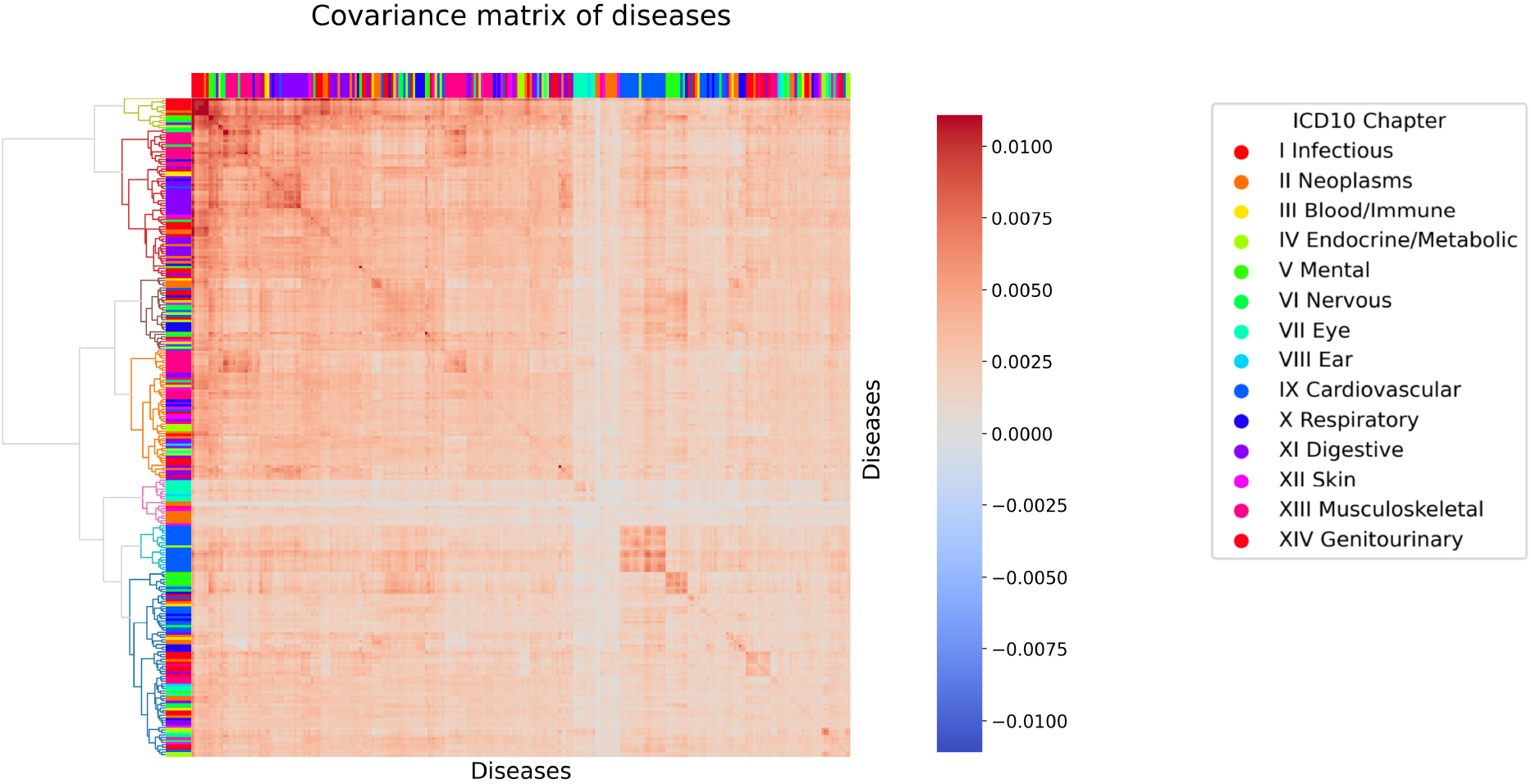
Covariance between diseases. Covariance **VV**^*⊤*^ between diseases. Diseases were clustered with hierarchical clustering using Ward’s method. Different clusters are color-coded on the dendrogram, diseases are color-coded by ICD-10 Chapter.

### A compact 10-protein panel for disease risk prediction

In clinical practice, measuring almost 3000 proteins for disease risk stratification can be infeasible. Therefore, we need other ways to predict patients’ disease risk. Instead of including the full panel with 3000 proteins, we can reduce the number of proteins needed by a greedy backwards selection approach, by systematically excluding proteins with lower variable importance. In this way, we can construct a compact panel including only 10 proteins, which is more suitable for clinical implementation. The panel retains *>* 99% of the performance of the full model.

Starting from the full panel with 3000 proteins, we select the 10-protein panel by recursively eliminating the protein with the lowest importance score calculated as

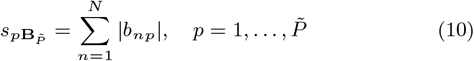

where 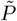 is the number of proteins included in a smaller panel and 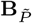 is the effect size matrix describing the influence of proteins on disease risk from the plasma proteins included in the smaller selection. Hyperparameter selection was repeated for each new selection of proteins, and MMAD-Risk was trained four times for the selected hyperparameter configuration. The next set of proteins was chosen based on the protein importance score from the model with the best performance on the validation set. Figure 5A shows the performance of MMAD-Risk depending on the number of proteins included in the analysis, panel (B) shows an overview of the selected proteins in the final panel and their functions. This small panel of proteins contains proteins that represent different functions, mostly related to immune and inflammatory signaling and metabolic and mitochondrial dysfunction. In addition, it also contains proteins associated with neurodegeneration, skin ageing and genomic and cellular stress.

**Figure 5.**
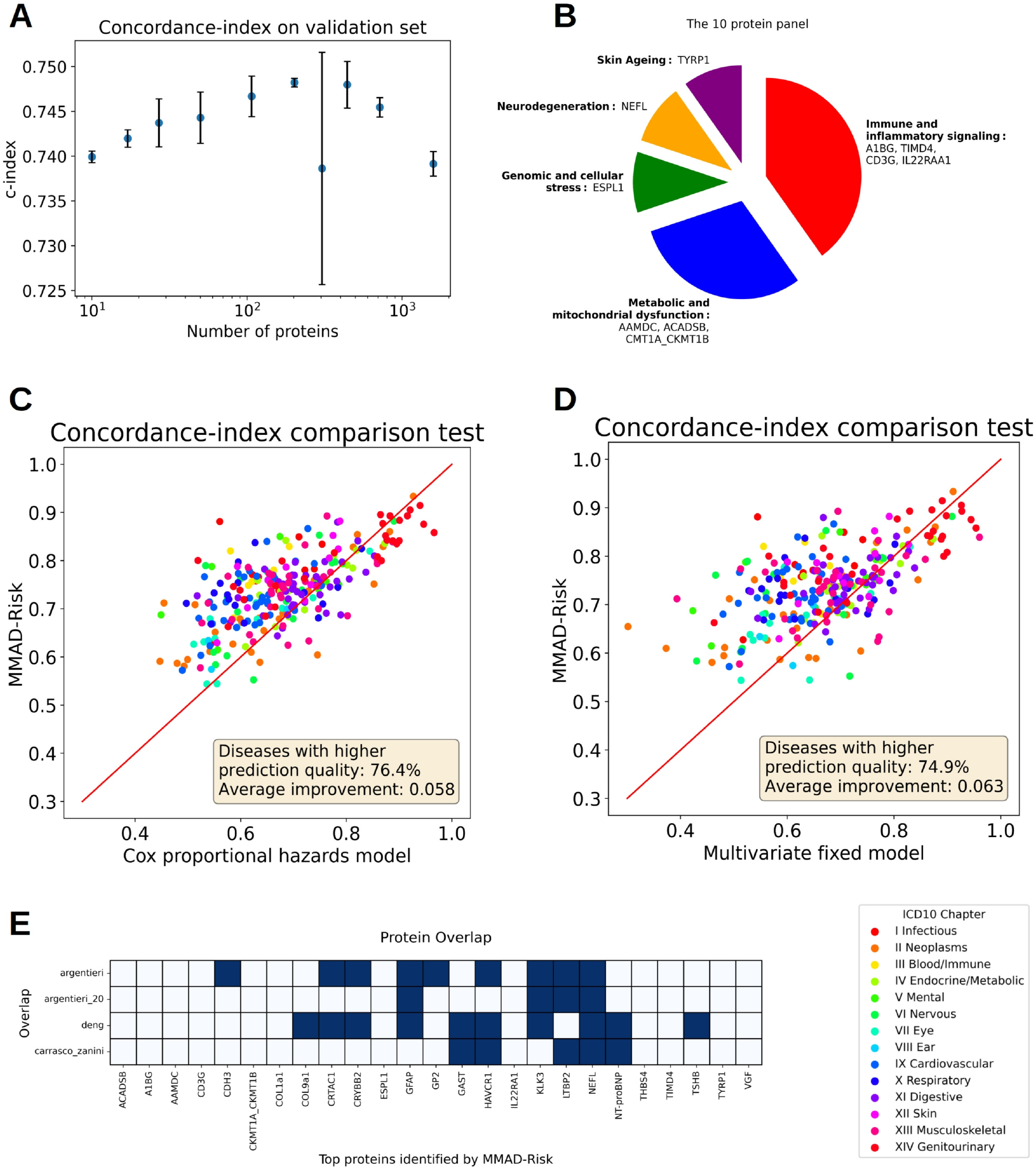
Ten-protein panel. (A) Performance of MMAD-Risk during greedy backwards selection based on different numbers of proteins included in the analysis. Error bars indicate standard deviation across four different runs. (B) Overview about the 10-protein panel and the function of these 10 proteins. (C) and (D) display model performance comparison on the 10-protein panel based on the c-index between MMAD-Risk and a Cox proportional hazards model and MMAD-Risk and a multivariate fixed survival analysis model, respectively. (E) Protein overlap between the top 3 proteins of each cluster, the 10-protein panel and proteins identified in major proteomic studies (Argentieri et al., 2024; Deng et al., 2025; Carrasco-Zanini et al., 2024).

We repeated the performance comparison of MMAD-Risk against a Cox proportional hazards model and the multivariate fixed effect model (Figure 5C,D). On the 10-protein panel, MMAD-Risk outperforms the competing models more substantially than on the full panel of 3000 proteins. MMAD-Risk outperforms the Cox model on 76.4% of diseases and improves the mean c-index by 0.058, whereas in comparison the multivariate fixed model, MMAD-Risk improves the disease risk prediction on 74.9% of diseases with a mean c-index improvement of 0.063.

Panel (E) shows the overlap of the three most important proteins per cluster and the 10-protein panel identified by MMAD-Risk with proteins identified by major proteomic studies from Argentieri et al. (2024), Deng et al. (2025) and Carrasco-Zanini et al. (2024). Out of the 25 proteins identified by MMAD-Risk, only 13 proteins overlap with at least one of these studies which is lower than expected.

Supplementary Figure 9 shows the influence of these 10 proteins on disease risk of all 271 diseases.

## Discussion

In this study, we present MMAD-Risk, a novel multivariate mixed survival analysis model based on variational inference. We demonstrate that this model outperforms more established survival analysis models on age-dependent disease risk prediction on the UK Biobank Pharma Proteomics Data Set. It is the first example of a multivariate mixed survival analysis model applied to hundreds of different outcomes, thereby overcoming previous technical limitations.

From a methodological point of view, we have developed an efficient method to include random effects to capture the correlations within a multivariate failure time response variable. To learn the model parameters, one cannot maximize the likelihood, as this involves an integral over the latent random effects variables (eq (2)). We therefore used variational inference to optimize the ELBO. We transfer the framework developed by Hui et al. (2017) and select the variational distribution by mirroring the structure of the posterior distribution in equation (2), as the product of a parameterless multivariate normal distribution *N*(**z**_*n*_|**0, I**_*K*_) and the product of univariate normals approximating the one-dimensional likelihoods of the responses. This parameter-efficient and simple form allowed us to decompose the complex multivariate problem into a series of tractable one-dimensional integrals. The analytical form of the ELBO and its gradients allowed us to train model parameters within minutes on tens of thousands of samples each with thousands of covariates and hundreds of outcomes. We are also exploring this variational ansatz in combination with stochastic variational inference to efficiently train models for structured additive distributional regression (Callegher et al., 2024).

We introduce a novel way of decomposing disease risks into factors of aging, allowing us to identify disease mechanisms shared across different organ systems. We propose to preprocess the protein plasma concentrations by calculating the deviation of the plasma protein concentration from the age-typical concentrations. This ensures that MMAD-Risk captures disease signatures instead of the proteomic signature of ageing.

Jiang et al. developed a multivariate survival analysis model based on deep learning (Jiang et al., 2025). However, their method does not implement a low-rank decomposition and cannot model random effects, and therefore MMAD-Risk is better suited when covariates cannot fully capture the comorbidity structure of covarying disease risks.

MMAD-Risk adds more evidence to the claim of Shmatko et al. (2025) that predicting the risk of getting several diseases together outperforms singe disease risk prediction for a lot of diseases. In addition to their model, MMAD-Risk adds insights on disease-related proteins and is easier to interpret since it is not based on deep learning.

In contrast to proteomic ageing clocks (Argentieri et al., 2024; Sedaghat et al., 2025) and proteomic organ ageing clocks (Oh et al., 2023, 2025; Wen, 2025; Wang et al., 2025a,b; Kivimäki et al., 2025), MMAD-Risk predicts disease risk directly instead of relying on a difference of a predicted proteomic and observed chronological age. The reliance of biological ageing clocks on the difference between predicted and chronological age, and the subsequent difficulty to distinguish statistical noise from biological signal has been criticized before (Ikram, 2024). The low rank matrix decomposition of **B** allows us to identify factors of ageing, i.e. different, distinct processes contributing to ageing. It does not need any preselection of proteins based on organ type. The clustering of **V** reveals that different ageing processes influence a lot of different types of disease, therefore an overly strict preselection of proteins might overlook the influence of less specific proteins on disease risk. In contrast, MMAD-Risk is able to identify shared disease mechanisms across different organ systems.

MMAD-Risk strongly outperforms Cox regression both on the full panel containing 3000 proteins and on the ten-protein panel. It only improves incrementally upon the multivariate fixed effect model on the full protein panel, but strongly outperforms it on the 10-protein data set. Our hypothesis is that much of the disease history can be deduced indirectly from the plasma protein concentrations of 3000 proteins, but when working with a small sample of proteins, the disease history becomes more informative.

We identified 10 clusters of diseases and the ten most predictive proteins in each of these clusters. In addition, we identify a small predictive set of only 10 proteins for which the model retains *>* 99% of its performance measured by the c-index on the test set. This minimal set can be useful in a clinical setting where only few proteins can be measured. Many of these proteins were not previously identified in major studies such as (Argentieri et al., 2024; Deng et al., 2025; Carrasco-Zanini et al., 2024). Therefore, more research is needed in order to assess the association between these proteins and age-dependent disease risk.

A limitation of our study is that it lacks validation on an external cohort such as FinnGen (Kurki et al., 2023) or the China Kadoorie Biobank (Chen et al., 2011). Additionally, it does not take the longitudinal development of plasma protein concentrations into account and relies on a single plasma protein measurement instead. Since the UK Biobank plasma proteomics data set only contains participants whose blood samples were drawn between ages 40 to 70, the results of this study cannot be transferred to other age groups. In this study, we used the Weibull survival function with a single shape parameter, which might not accurately describe survival time distribution for some diseases.

MMAD-Risk is the first multivariate mixed survival method to be applied to hundreds of different response variables. By transferring a variational inference approach where the ELBO can be calculated analytically (Hui et al., 2017) to survival analysis, we can optimize MMAD-Risk fast and efficiently. In contrast to the R implementation in gllvm by Hui et al. (2017), which shows convergence issues on real data, MMAD-Risk converges reliably. We demonstrated that the mixed modeling approach is especially useful in situations where only few external covariates and a high number of correlated response variables are available.

In this study, we evaluated the potential of MMAD-Risk for age-dependent disease risk prediction. However, multivariate mixed survival analysis holds promise beyond biomedicine. Examples are industrial machine maintenance, where component failures are often correlated, finance, where systemic market crises can co-occur, and climate science, where natural catastrophes are driven by shared latent environmental forces.

## Author Contributions

A.M.H. designed the research, developed the software, performed the analyses, and wrote the paper; J.S. supervised the project, designed the research, and wrote the paper.

## Ethics Statement

This study is based entirely on computational analysis of publicly available data from UK Biobank. No ethical approval was required as no human subjects, human tissue, or animals were involved.

## Data availability

This research has been conducted using the UK Biobank Resource under application number 532367. The UK Biobank data can be accessed by a procedure described in https://www.ukbiobank.ac.uk/use-our-data/.

## Code availability

The source code for MMAD-Risk is available on GitHub https://github.com/soedinglab/MMAD-Risk repository under the Apache License 2.0.

## Acknowledgments

We thank Thomas Kneib and Anne-Christin Hauschild for discussions and the International Max Planck Research School for Genome Science and especially Henriette Irmer for supporting AMH’s dissertation.

During the preparation of this work, AMH used Qwen-3-30B-A3B-Instruct 2507 and Claude Sonnet 4.6 and 4.7 to assist debugging the software and proofreading and to suggest rephrasing selected sentences. The authors reviewed, tested, and optimized all generated code independently and take full responsibility for the accuracy of the computational results.

