## Supplementary Material for "MMAD-Risk: Multivariate Mixed Survival Analysis for the Prediction of Age-Dependent Disease Risks from Plasma Proteomes"

### Full deviation of the Evidence Lower Bound

We present the full deviation of the ELBO (evidence lower bound) from the methods part of the main article.

We have selected the following variational distribution:

$$q(\mathbf{z}_n|\mu_n,\lambda_n) \propto \prod_{d=1}^{D} \mathscr{N}\left(\mathbf{v}_d^\top \mathbf{z}_n + \mathbf{b}_d^\top \mathbf{x}_n|\mu_{nd},\lambda_{nd}^{-1}\right) \times \mathscr{N}(\mathbf{z}_n|0,\mathbf{I}_K).$$

By completing the square, we can show that the closed from of the log likelihood of the variational distribution is given by

$$\begin{aligned}
\ln q(\mathbf{z}_n|\mu_n,\operatorname{diag}(\lambda_{nd})) &= \sum_{d=1}^{D} \ln \mathscr{N}\left(\mathbf{v}_d^\top \mathbf{z}_n + \mathbf{b}_d^\top \mathbf{x}_n|\mu_{nd},\lambda_{nd}^{-1}\right) + \ln \mathscr{N}(\mathbf{z}_n|0,\mathbf{I}_K) + \text{const} \\
&= \sum_{d=1}^{D} \ln \mathscr{N}\left(\mathbf{v}_d^\top z_{nd}|\nu_{nd},\lambda_{nd}^{-1}\right) + \ln \mathscr{N}(\mathbf{z}_n|0,\mathbf{I}_K) + \text{const} \\
&= -\frac{1}{2}\sum_{d=1}^{D} \lambda_{nd}\left(\mathbf{v}_d^\top z_{nd} - \nu_{nd}\right)^2 - \frac{1}{2}\mathbf{z}_n^\top \mathbf{I}_K \mathbf{z}_n + \text{const} \\
&= -\frac{1}{2}\sum_{d=1}^{D} \left(\mathbf{z}_n^\top \mathbf{v}_d - \nu_{nd}\right)\lambda_{nd}\left(\mathbf{v}_d^\top \mathbf{z}_n - \nu_{nd}\right) - \frac{1}{2}\mathbf{z}_n^\top \mathbf{I}_K \mathbf{z}_n + \text{const} \\
&= -\frac{1}{2}\left(\mathbf{z}_n^\top \mathbf{V}^\top \operatorname{diag}(\lambda_{nd})\mathbf{V}\mathbf{z}_n - 2\mathbf{z}_n^\top \mathbf{V}^\top \operatorname{diag}(\lambda_{nd})\nu_n + \mathbf{z}_n^\top \mathbf{I}_K \mathbf{z}_n\right) + \text{const} \\
&= -\frac{1}{2}\left(\mathbf{z}_n^\top (\mathbf{V}^\top \operatorname{diag}(\lambda_{nd})\mathbf{V} + \mathbf{I}_K)\mathbf{z}_n - 2\mathbf{z}_n^\top \mathbf{V}^\top \operatorname{diag}(\lambda_{nd})\nu_n\right) + \text{const} \\
&= \ln \mathscr{N}(\mathbf{z}_n|\mathbf{m}_n,\mathbf{S}_n)\,, \qquad \text{(S1)}
\end{aligned}$$

where

$$\mathbf{S}_n = (\mathbf{V}^\top \operatorname{diag}(\lambda_{nd})\mathbf{V} + \mathbf{I}_K)^{-1} \in \mathbb{R}^{K\times K}, \qquad \text{(S2)}$$

$$\nu_n = \mu_n - \mathbf{B}\mathbf{x}_n \in \mathbb{R}^D,$$

$$\mathbf{m}_n = \mathbf{S}_n \mathbf{V}^\top \operatorname{diag}(\lambda_{nd})\nu_n \in \mathbb{R}^K. \qquad \text{(S3)}$$

Based on these results, we can decompose the ELBO into the three following terms:

$$\begin{aligned}
\mathrm{ELBO}(\phi,\theta) &= \sum_{n=1}^{N} \mathbb{E}_{q(\mathbf{z}_n|\mathbf{m}_n,\mathbf{S}_n)}\left[\ln \frac{p(\mathbf{y}_n,\mathbf{z}_n|\mathbf{x}_n,\delta_n,\theta)}{q(\mathbf{z}_n|\phi)}\right] \\
&= \sum_{n=1}^{N} \mathbb{E}_{q(\mathbf{z}_n|\mathbf{m}_n,\mathbf{S}_n)}\left[\ln \frac{\prod_{d=1}^{D} p(\mathbf{y}_{nd}|\theta,\delta_n,\mathbf{x}_n)\,\mathcal{N}(\mathbf{z}_n|0,\mathbf{I}_K)}{\mathcal{N}(\mathbf{z}_n|\mathbf{m}_n,\mathbf{S}_n)}\right] \\
&= \sum_{n=1}^{N} \Big(\mathbb{E}_{q(\mathbf{z}_n|\mathbf{m}_n,\mathbf{S}_n)}\big[\ln \mathcal{N}(\mathbf{z}_n|0,\mathbf{I}_K)\big] \\
&\quad - \mathbb{E}_{q(\mathbf{z}_n|\mathbf{m}_n,\mathbf{S}_n)}\big[\ln \mathcal{N}(\mathbf{z}_n|\mathbf{m}_n,\mathbf{S}_n)\big] \\
&\quad + \sum_{d=1}^{D} \mathbb{E}_{q(\mathbf{z}_n|\mathbf{m}_n,\mathbf{S}_n)}\big[\ln p(y_{nd}|\mathbf{x}_n,\delta_{nd},\theta)\big]\Big)\,. \qquad \text{(S4)}
\end{aligned}$$

These three terms can be further transformed to

$$\begin{aligned}
&\mathbb{E}_{q(\mathbf{z}_n|\mathbf{m}_n,\mathbf{S}_n)}\big[\ln \mathcal{N}(\mathbf{z}_n|0,\mathbf{I}_K)\big] = -\frac{1}{2}\Big(\mathrm{Tr}(\mathbf{S}_n) + \mathbf{m}_n^\top \mathbf{m}_n\Big) + \mathrm{const} \\
&\mathbb{E}_{q(\mathbf{z}_n|\mathbf{m}_n,\mathbf{S}_n)}\big[\ln \mathcal{N}(\mathbf{z}_n|\mathbf{m}_n,\mathbf{S}_n)\big] = -\frac{1}{2}\ln\det \mathbf{S}_n + \mathrm{const}\,, \\
&\mathbb{E}_{q(\mathbf{z}_n|\mathbf{m}_n,\mathbf{S}_n)}\big[\ln p(\mathbf{y}_n|\mathbf{x}_n,\delta_n,\theta)\big] \\
&\quad = \sum_{d=1}^{D} \mathbb{E}_{q(\mathbf{z}_n|\mathbf{m}_n,\mathbf{S}_n)}\big[\ln p(y_{nd}|\mathbf{x}_n,\delta_{nd},\theta)\big] \\
&\quad = \sum_{d=1}^{D} \mathbb{E}_{\mathcal{N}(\eta_{nd}|\mathbf{b}_d^\top \mathbf{x}_n + \mathbf{v}_d^\top \mathbf{m}_n,\, \mathbf{v}_d^\top \mathbf{S}_n \mathbf{v}_d)}\big[\ln p(\eta_{nd}|\mathbf{x}_n,\delta_{nd},\theta)\big] + \mathrm{const} \\
&\quad = \sum_{d=1}^{D} \mathbb{E}_{\mathcal{N}(\eta_{nd}|\mathbf{b}_d^\top \mathbf{x}_n + \mathbf{v}_d^\top \mathbf{m}_n,\, \mathbf{v}_d^\top \mathbf{S}_n \mathbf{v}_d)}\left[\delta_{nd}\left(\ln \frac{\kappa_d y_{nd}^{\kappa_d - 1}}{\tau^{\kappa_d}} + \kappa_d \eta_{nd}\right) - \frac{y_{nd}^{\kappa_d}}{\tau^{\kappa_d}} e^{\kappa_d \eta_{nd}}\right] + \mathrm{const} \\
&\quad = \sum_{d=1}^{D} \left[\delta_{nd}\left(\ln \frac{\kappa_d y_{nd}^{\kappa_d - 1}}{\tau^{\kappa_d}} + \kappa_d(\mathbf{b}_d^\top \mathbf{x}_n + \mathbf{v}_d^\top \mathbf{m}_n)\right)\right. \\
&\qquad \left. - \frac{y_{nd}^{\kappa_d}}{\tau^{\kappa_d}} \exp\left(\frac{\kappa_d^2}{2}\mathbf{v}_d^\top \mathbf{S}_n \mathbf{v}_d + \kappa_d(\mathbf{b}_d^\top \mathbf{x}_n + \mathbf{v}_d^\top \mathbf{m}_n)\right)\right] + \mathrm{const.}
\end{aligned}$$

### Masking diagnoses made before time of sample collection

We aim to predict disease risk from UK Biobank participants' plasma proteomes. In order to capture the signature of future disease and to avoid reverse causation, we introduce a mask $\omega \in \mathbb{R}^{n\times d}$ that indicates whether a disease was diagnosed before or after the blood sample collection from UK Biobank participants:

$$\begin{aligned}
\omega_{nd} &= 0 \quad \text{diagnosis before blood sample collection}\,, \\
\omega_{nd} &= 1 \quad \text{otherwise}\,.
\end{aligned}$$

We add it to the third term of the ELBO since this is the only term where the disease status of UK Biobank participants enters:

$$\begin{aligned}
&\mathbb{E}_{q(\mathbf{z}_n|\mathbf{m}_n,\mathbf{S}_n)}\left[\ln p(\mathbf{y}_n|\mathbf{x}_n,\delta_n,\theta)\right] \\
&= \sum_{d=1}^{D} \mathbb{E}_{q(\mathbf{z}_n|\mathbf{m}_n,\mathbf{S}_n)}\left[\ln p(y_{nd}|\mathbf{x}_n,\delta_{nd},\theta)\right] \\
&= \sum_{d=1}^{D} \omega_{nd}\Bigg[\delta_{nd}\left(\ln\frac{\kappa_d y_{nd}^{\kappa_d-1}}{\tau^{\kappa_d}} + \kappa_d(\mathbf{b}_d^\top\mathbf{x}_n + \mathbf{v}_d^\top\mathbf{m}_n)\right) \\
&\quad - \frac{y_{nd}^{\kappa_d}}{\tau^{\kappa_d}}\exp\left(\frac{\kappa_d^2}{2}\mathbf{v}_d^\top\mathbf{S}_n\mathbf{v}_d + \kappa_d(\mathbf{b}_d^\top\mathbf{x}_n + \mathbf{v}_d^\top\mathbf{m}_n)\right)\Bigg] + \text{const.}
\end{aligned}$$

This allows us to mask patients who were diagnosed before the blood sample collection and prevents us from accidentally picking up the signature of a medication that a diagnosed patient receives for their disease without having to exclude these patients from the analysis.

### Data Preprocessing

Plots of two examples of the calculation of the age-dependent normalized protein concentration, Col9a1 and KLK3, are shown in Figures 1 and 2. For the spline regression, we selected third degree splines with six nodes and periodic extrapolation.

### Regularization

Since not all proteins are expected to have an effect on ageing and age-dependent disease risks, we place a group lasso prior on $\mathbf{B}$:

$$-\lambda_G \sum_{p=1}^{P}\left(\sum_{d=1}^{D} b_{dp}^2\right)^{1/2},$$

where $P$ is the number of proteins included into the model. Additionally, we place a ridge regression penalty on $\mathbf{B}$ and a lasso penalty on $\mathbf{V}$. The regularization strength was determined by the model performance on the validation set. The selected regularization strengths were $\lambda_G = 30$ for the group lasso, $\lambda_r = 20$ for the ridge penalty and $\lambda_V = 30$ for the lasso penalty via the performance on the validation set during a grid search.

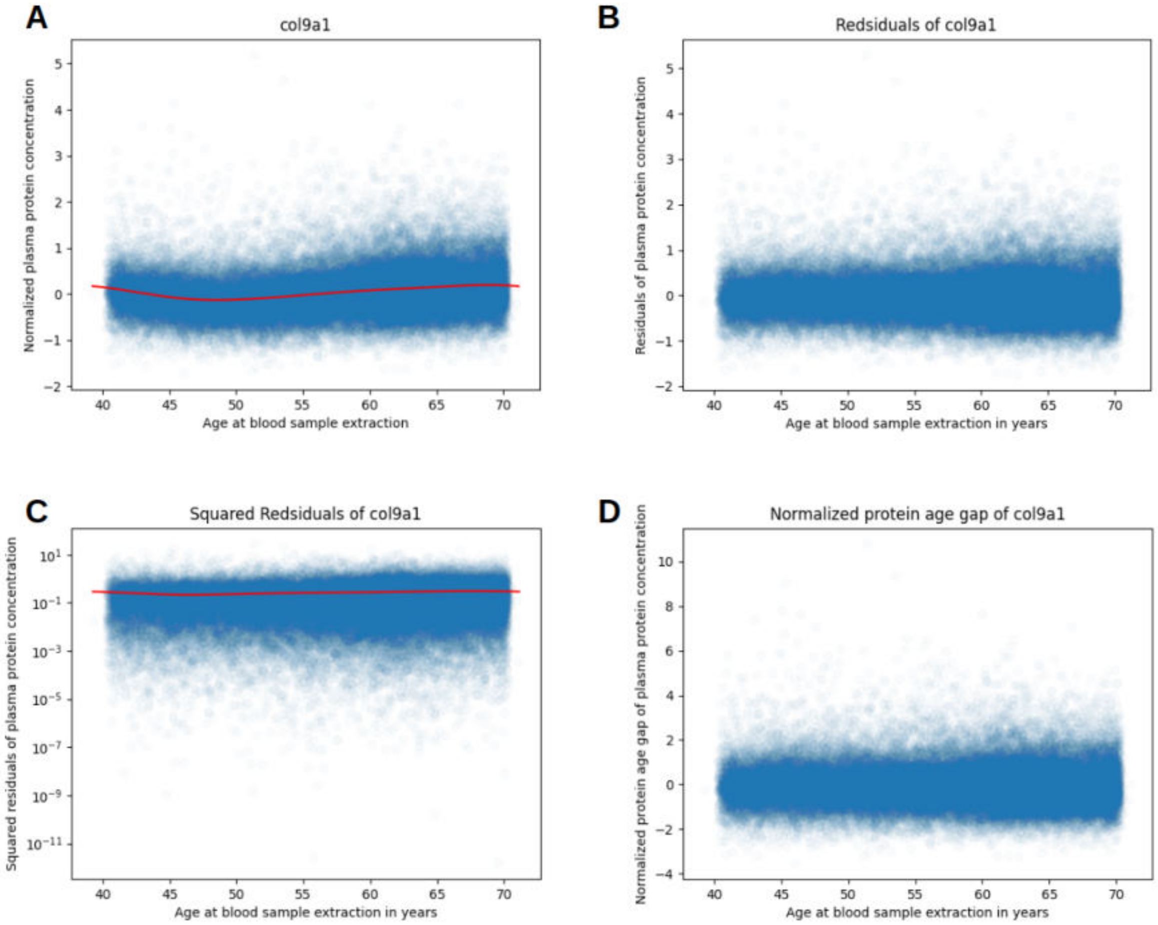


Figure 1: This figure shows the protein concentration preprocessing step of COL9a1. (A): Protein Concentration with spline regression, (B): residuals of the spline regression, (C): Squared residuals fitted with another spline regression, (D): Age-dependent normalized protein concentration.

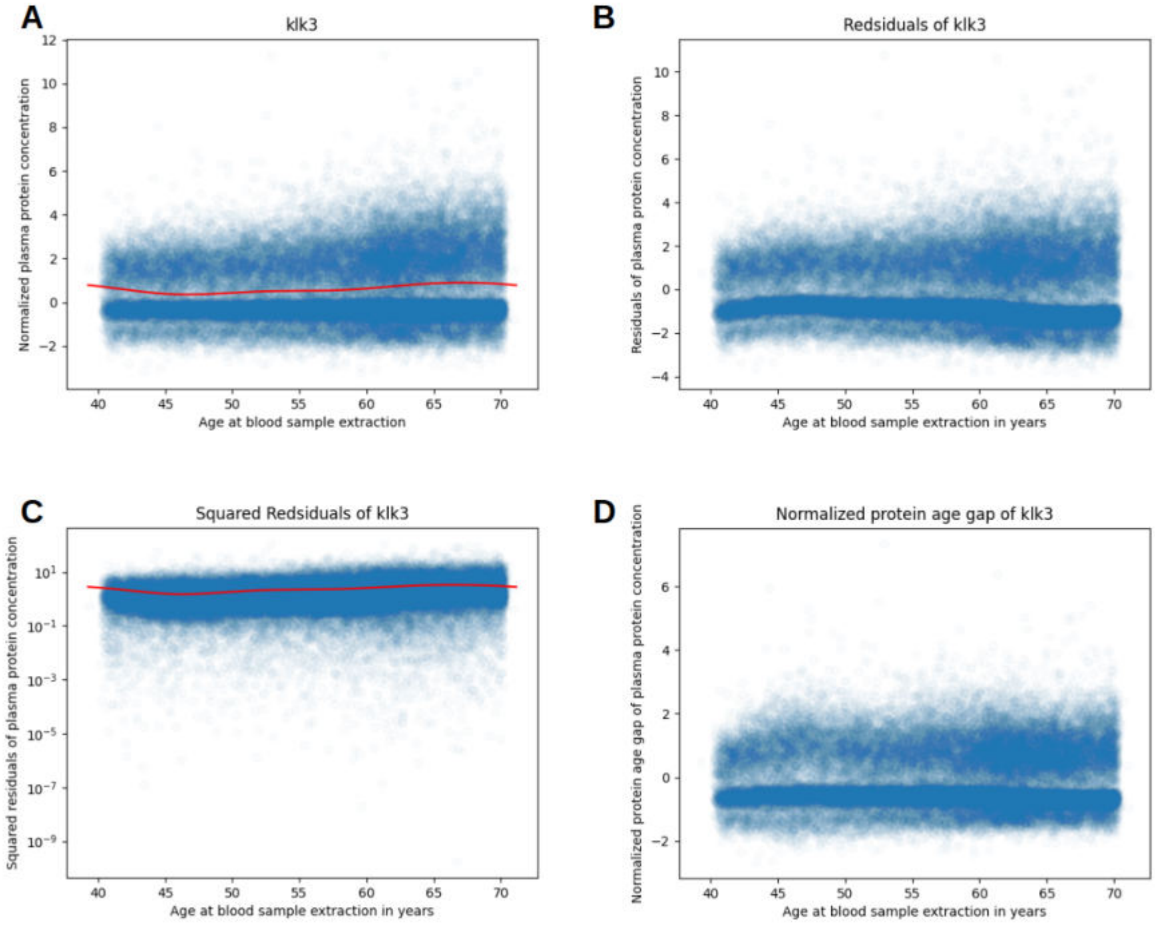


Figure 2: This figure shows the protein concentration preprocessing step of KLK3. (A): Protein Concentration with spline regression, (B): residuals of the spline regression, (C): Squared residuals fitted with another spline regression, (D): Age-dependent normalized protein concentration.

### Selection of $K$

In order to identify the optimal $K$, i.e. the rank of the matrices $\mathbf{B}, \mathbf{V}$ and $\mathbf{W}$, we trained MMAD-Risk four times on the data set with 3000 proteins and evaluated the performance of MMAD-risk based on the c-index. The results can be seen in Figure 3. Based on these results, we chose $K = 15$ since this achieves the optimal performance while requiring fewer parameters than for $K = 25$ or $K = 30$.

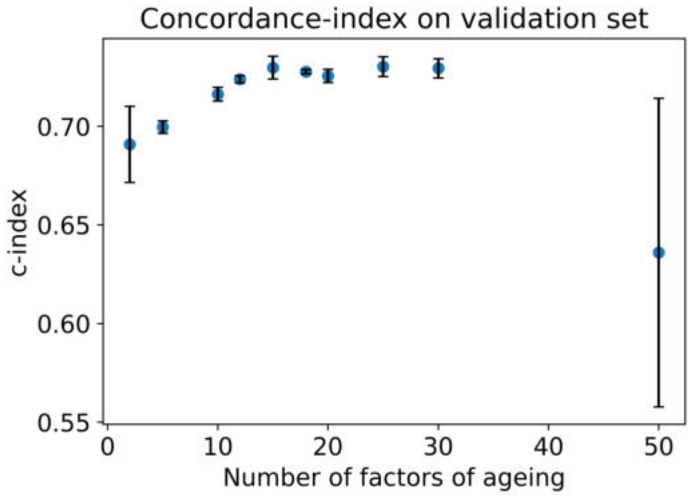


Figure 3: Plot of c-index on the validation data set dependent on the number of factors ageing $K$. For each $K$, the model was run four times. Dots represent the mean and error bars the standard deviation across all four runs.

### Comparison of MMAD-Risk with a univariate mixed model and logistic regression

In addition to the comparisons of MMAD-Risk to a multivariate fixed survival analysis model and Cox regression, we compare our model to a univariate mixed survival analysis model and logistic regression. The log likelihood of the univariate mixed model is similar to the log likelihood, but its effect size matrix $\mathbf{B}$ is not decomposed into two lower-rank matrices and the random effects $\mathbf{V}\mathbf{z}_n$ are one dimensional.

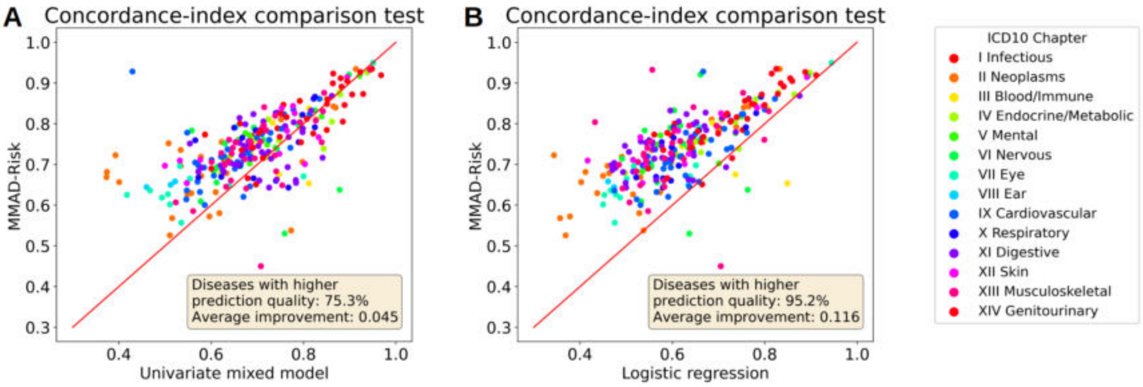


Figure 4: Panels (A) and (B) display model performance comparison based on the c-index between MMAD-Risk and a univariate mixed survival analysis model and MMAD-Risk and logistic regression, respectively.

### Clustering of $\mathbf{V}$

Before clustering, we first divide every column of $\mathbf{V}$ by its column sum: $\mathbf{v}_k^\dagger = \mathbf{v}_k / \sum_{d=1}^{D} v_{dk}$ for $k = 1, \ldots, K$, where $\mathbf{v}_1, \ldots, \mathbf{v}_K$ are the columns of $\mathbf{V}$ and $\tilde{\mathbf{V}} = [\tilde{\mathbf{v}}_1, \ldots, \tilde{\mathbf{v}}_K]$ is a column-normalized matrix. In the next step, the rows of $\mathbf{V}$ are divided by their standard deviation: $\tilde{\mathbf{v}}_d = \mathbf{v}_d^\dagger / \mathrm{std}(\mathbf{v}_d^\dagger)$ for $d = 1, \ldots, D$, where $\mathbf{v}_1^\dagger, \ldots, \mathbf{v}_D^\dagger$ are the rows of $\mathbf{V}^\dagger$. The rows and columns of $\tilde{\mathbf{V}}$ then are clustered using Ward's method.

The matrix $\mathbf{V}$ was clustered using Ward's method for agglomerative clustering and the number is clusters was set to 10 based on the elbow method. The corresponding elbow plot can be seen in Figure 5.

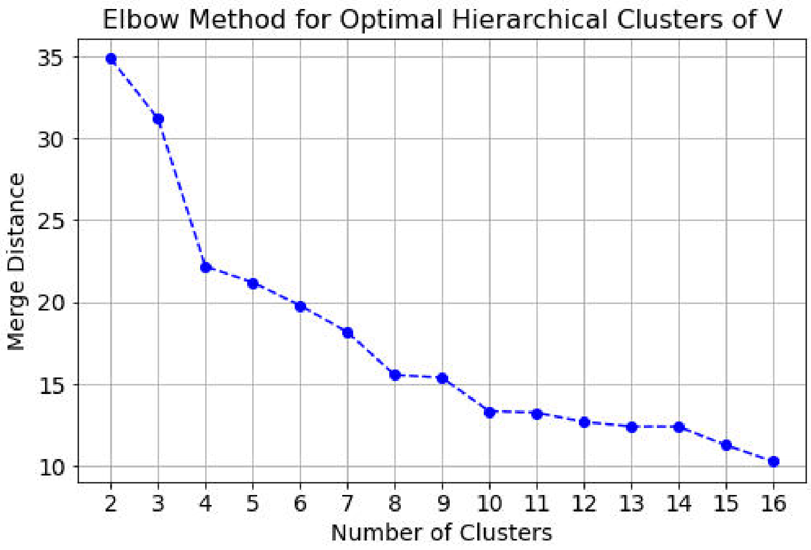


Figure 5: Elbow plot for the clusters of $\mathbf{V}$.

The full dendrogram with disease labels from Figure 3A is shown in Figure 6:

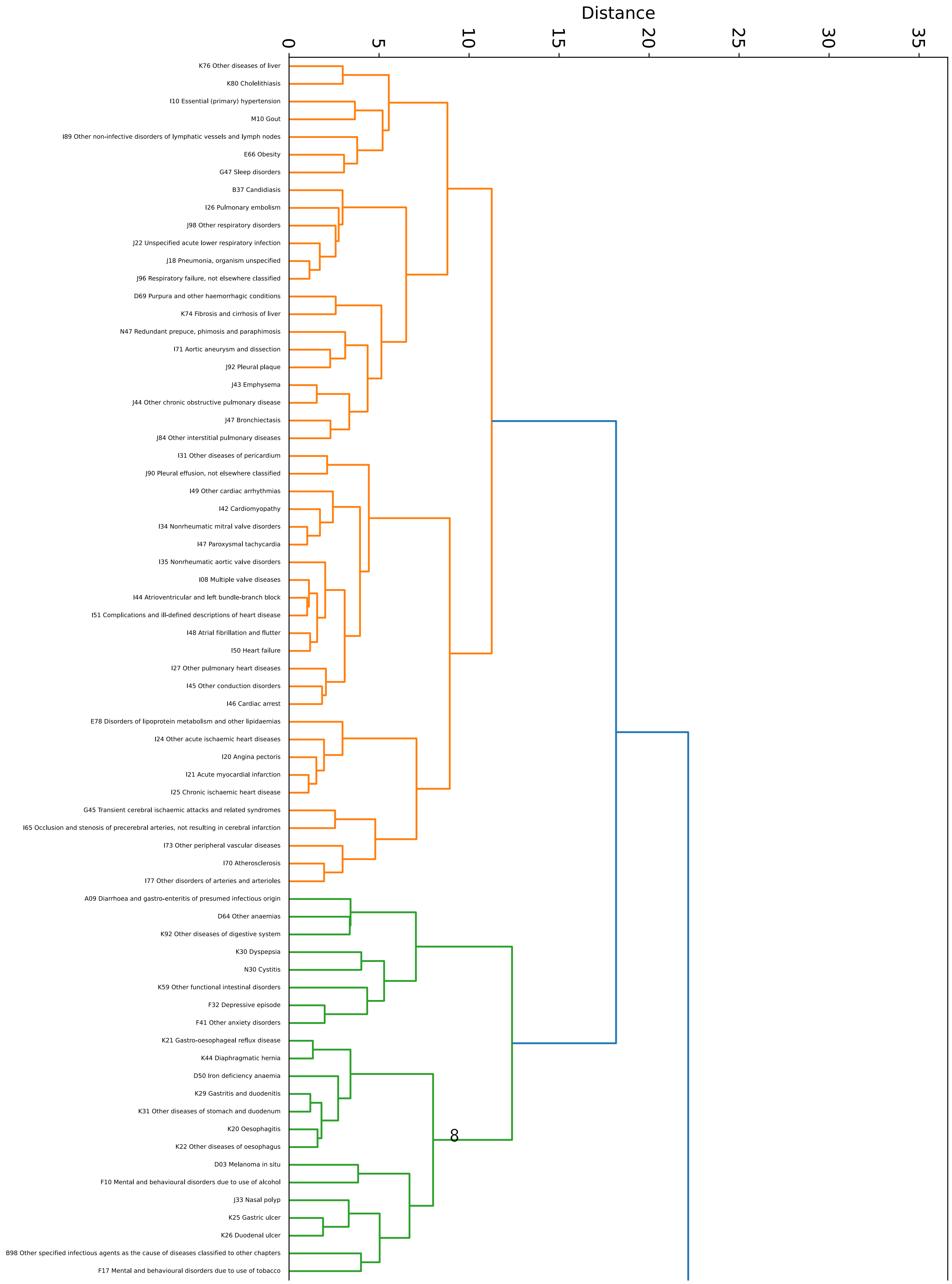
Distance
0
5
10
15
20
25
30
35
K76 Other diseases of liver
K80 Cholelithiasis
I10 Essential (primary) hypertension
M10 Gout
I89 Other non-infective disorders of lymphatic vessels and lymph nodes
E66 Obesity
G47 Sleep disorders
B37 Candidiasis
I26 Pulmonary embolism
J98 Other respiratory disorders
J22 Unspecified acute lower respiratory infection
J18 Pneumonia, organism unspecified
J96 Respiratory failure, not elsewhere classified
D69 Purpura and other haemorrhagic conditions
K74 Fibrosis and cirrhosis of liver
N47 Redundant prepuce, phimosis and paraphimosis
I71 Aortic aneurysm and dissection
J92 Pleural plaque
J43 Emphysema
J44 Other chronic obstructive pulmonary disease
J47 Bronchiectasis
J84 Other interstitial pulmonary diseases
I31 Other diseases of pericardium
J90 Pleural effusion, not elsewhere classified
I49 Other cardiac arrhythmias
I42 Cardiomyopathy
I34 Nonrheumatic mitral valve disorders
I47 Paroxysmal tachycardia
I35 Nonrheumatic aortic valve disorders
I08 Multiple valve diseases
I44 Atrioventricular and left bundle-branch block
I51 Complications and ill-defined descriptions of heart disease
I48 Atrial fibrillation and flutter
I50 Heart failure
I27 Other pulmonary heart diseases
I45 Other conduction disorders
I46 Cardiac arrest
E78 Disorders of lipoprotein metabolism and other lipidaemias
I24 Other acute ischaemic heart diseases
I20 Angina pectoris
I21 Acute myocardial infarction
I25 Chronic ischaemic heart disease
G45 Transient cerebral ischaemic attacks and related syndromes
I65 Occlusion and stenosis of precerebral arteries, not resulting in cerebral infarction
I73 Other peripheral vascular diseases
I70 Atherosclerosis
I77 Other disorders of arteries and arterioles
A09 Diarrhoea and gastro-enteritis of presumed infectious origin
D64 Other anaemias
K92 Other diseases of digestive system
K30 Dyspepsia
N30 Cystitis
K59 Other functional intestinal disorders
F32 Depressive episode
F41 Other anxiety disorders
K21 Gastro-oesophageal reflux disease
K44 Diaphragmatic hernia
D50 Iron deficiency anaemia
K29 Gastritis and duodenitis
K31 Other diseases of stomach and duodenum
K20 Oesophagitis
K22 Other diseases of oesophagus
D03 Melanoma in situ
F10 Mental and behavioural disorders due to use of alcohol
J33 Nasal polyp
K25 Gastric ulcer
K26 Duodenal ulcer
B98 Other specified infectious agents as the cause of diseases classified to other chapters
F17 Mental and behavioural disorders due to use of tobacco

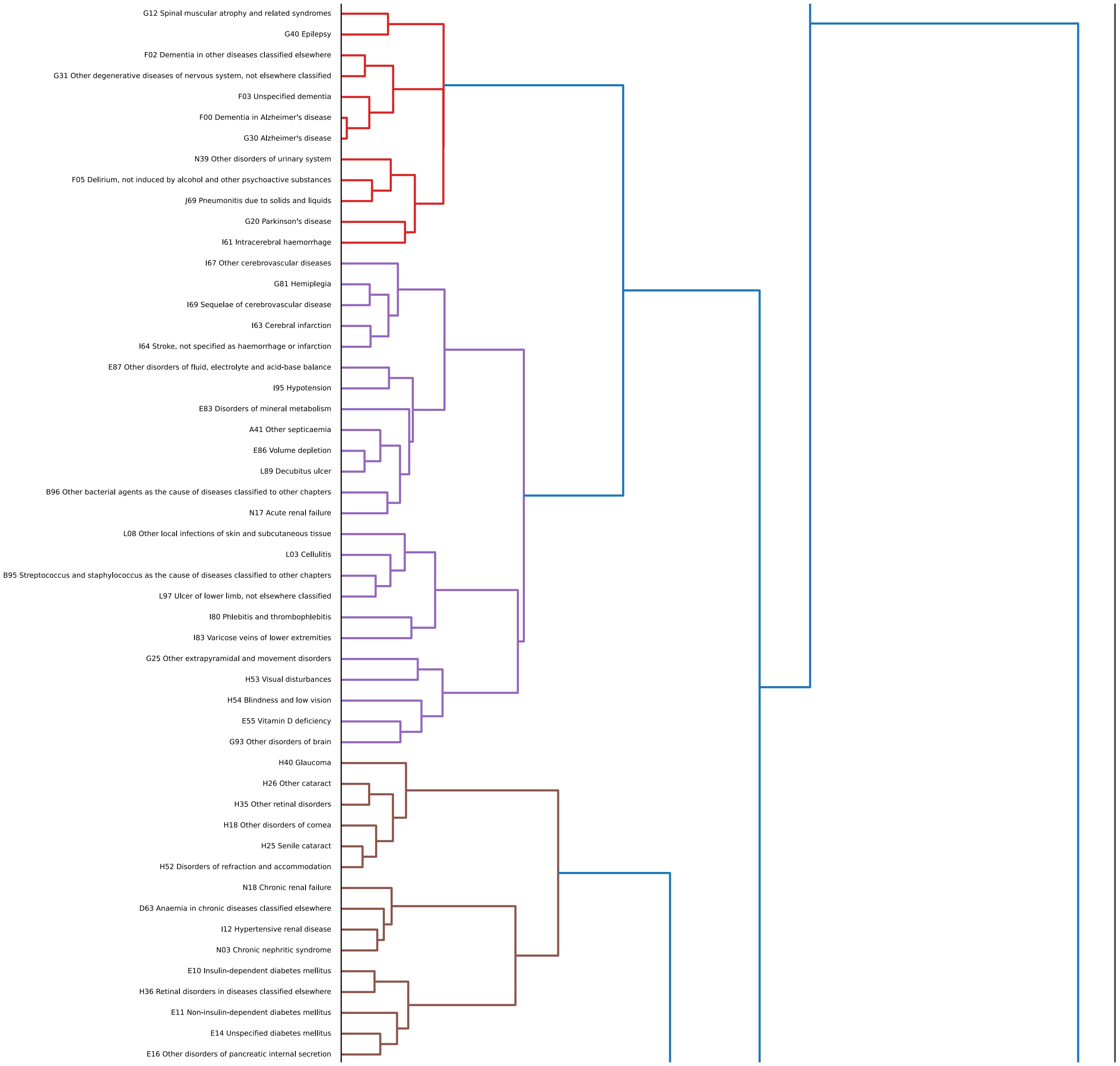
G12 Spinal muscular atrophy and related syndromes
G40 Epilepsy
F02 Dementia in other diseases classified elsewhere
G31 Other degenerative diseases of nervous system, not elsewhere classified
F03 Unspecified dementia
F00 Dementia in Alzheimer's disease
G30 Alzheimer's disease
N39 Other disorders of urinary system
F05 Delirium, not induced by alcohol and other psychoactive substances
J69 Pneumonitis due to solids and liquids
G20 Parkinson's disease
I61 Intracerebral haemorrhage
I67 Other cerebrovascular diseases
G81 Hemiplegia
I69 Sequelae of cerebrovascular disease
I63 Cerebral infarction
I64 Stroke, not specified as haemorrhage or infarction
E87 Other disorders of fluid, electrolyte and acid-base balance
I95 Hypotension
E83 Disorders of mineral metabolism
A41 Other septicaemia
E86 Volume depletion
L89 Decubitus ulcer
B96 Other bacterial agents as the cause of diseases classified to other chapters
N17 Acute renal failure
L08 Other local infections of skin and subcutaneous tissue
L03 Cellulitis
B95 Streptococcus and staphylococcus as the cause of diseases classified to other chapters
L97 Ulcer of lower limb, not elsewhere classified
I80 Phlebitis and thrombophlebitis
I83 Varicose veins of lower extremities
G25 Other extrapyramidal and movement disorders
H53 Visual disturbances
H54 Blindness and low vision
E55 Vitamin D deficiency
G93 Other disorders of brain
H40 Glaucoma
H26 Other cataract
H35 Other retinal disorders
H18 Other disorders of cornea
H25 Senile cataract
H52 Disorders of refraction and accommodation
N18 Chronic renal failure
D63 Anaemia in chronic diseases classified elsewhere
I12 Hypertensive renal disease
N03 Chronic nephritic syndrome
E10 Insulin-dependent diabetes mellitus
H36 Retinal disorders in diseases classified elsewhere
E11 Non-insulin-dependent diabetes mellitus
E14 Unspecified diabetes mellitus
E16 Other disorders of pancreatic internal secretion

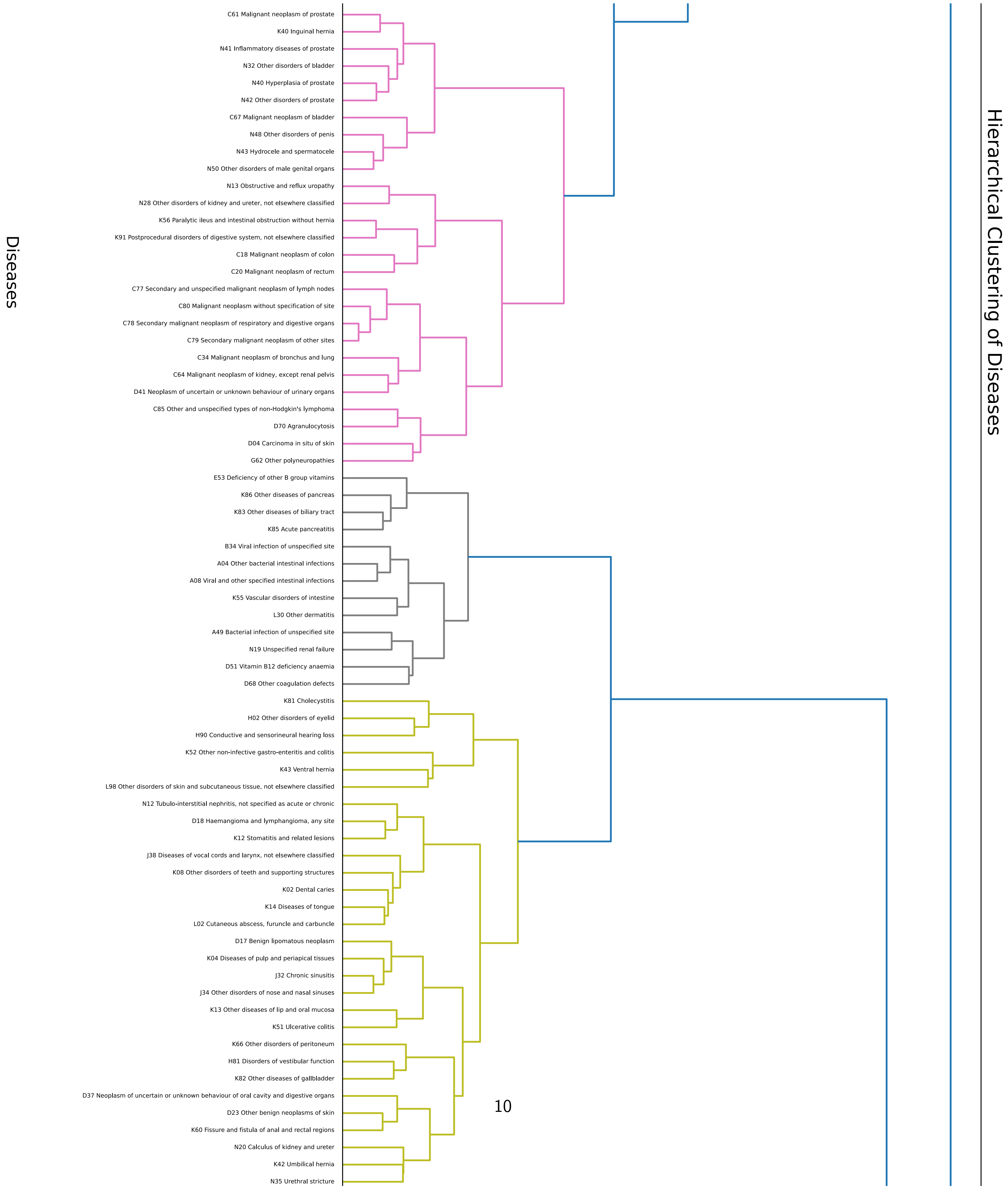
Hierarchical Clustering of Diseases
Diseases
C61 Malignant neoplasm of prostate
K40 Inguinal hernia
N41 Inflammatory diseases of prostate
N32 Other disorders of bladder
N40 Hyperplasia of prostate
N42 Other disorders of prostate
C67 Malignant neoplasm of bladder
N48 Other disorders of penis
N43 Hydrocele and spermatocele
N50 Other disorders of male genital organs
N13 Obstructive and reflux uropathy
N28 Other disorders of kidney and ureter, not elsewhere classified
K56 Paralytic ileus and intestinal obstruction without hernia
K91 Postprocedural disorders of digestive system, not elsewhere classified
C18 Malignant neoplasm of colon
C20 Malignant neoplasm of rectum
C77 Secondary and unspecified malignant neoplasm of lymph nodes
C80 Malignant neoplasm without specification of site
C78 Secondary malignant neoplasm of respiratory and digestive organs
C79 Secondary malignant neoplasm of other sites
C34 Malignant neoplasm of bronchus and lung
C64 Malignant neoplasm of kidney, except renal pelvis
D41 Neoplasm of uncertain or unknown behaviour of urinary organs
C85 Other and unspecified types of non-Hodgkin's lymphoma
D70 Agranulocytosis
D04 Carcinoma in situ of skin
G62 Other polyneuropathies
E53 Deficiency of other B group vitamins
K86 Other diseases of pancreas
K83 Other diseases of biliary tract
K85 Acute pancreatitis
B34 Viral infection of unspecified site
A04 Other bacterial intestinal infections
A08 Viral and other specified intestinal infections
K55 Vascular disorders of intestine
L30 Other dermatitis
A49 Bacterial infection of unspecified site
N19 Unspecified renal failure
D51 Vitamin B12 deficiency anaemia
D68 Other coagulation defects
K81 Cholecystitis
H02 Other disorders of eyelid
H90 Conductive and sensorineural hearing loss
K52 Other non-infective gastro-enteritis and colitis
K43 Ventral hernia
L98 Other disorders of skin and subcutaneous tissue, not elsewhere classified
N12 Tubulo-interstitial nephritis, not specified as acute or chronic
D18 Haemangioma and lymphangioma, any site
K12 Stomatitis and related lesions
J38 Diseases of vocal cords and larynx, not elsewhere classified
K08 Other disorders of teeth and supporting structures
K02 Dental caries
K14 Diseases of tongue
L02 Cutaneous abscess, furuncle and carbuncle
D17 Benign lipomatous neoplasm
K04 Diseases of pulp and periapical tissues
J32 Chronic sinusitis
J34 Other disorders of nose and nasal sinuses
K13 Other diseases of lip and oral mucosa
K51 Ulcerative colitis
K66 Other disorders of peritoneum
H81 Disorders of vestibular function
K82 Other diseases of gallbladder
D37 Neoplasm of uncertain or unknown behaviour of oral cavity and digestive organs
D23 Other benign neoplasms of skin
K60 Fissure and fistula of anal and rectal regions
N20 Calculus of kidney and ureter
K42 Umbilical hernia
N35 Urethral stricture

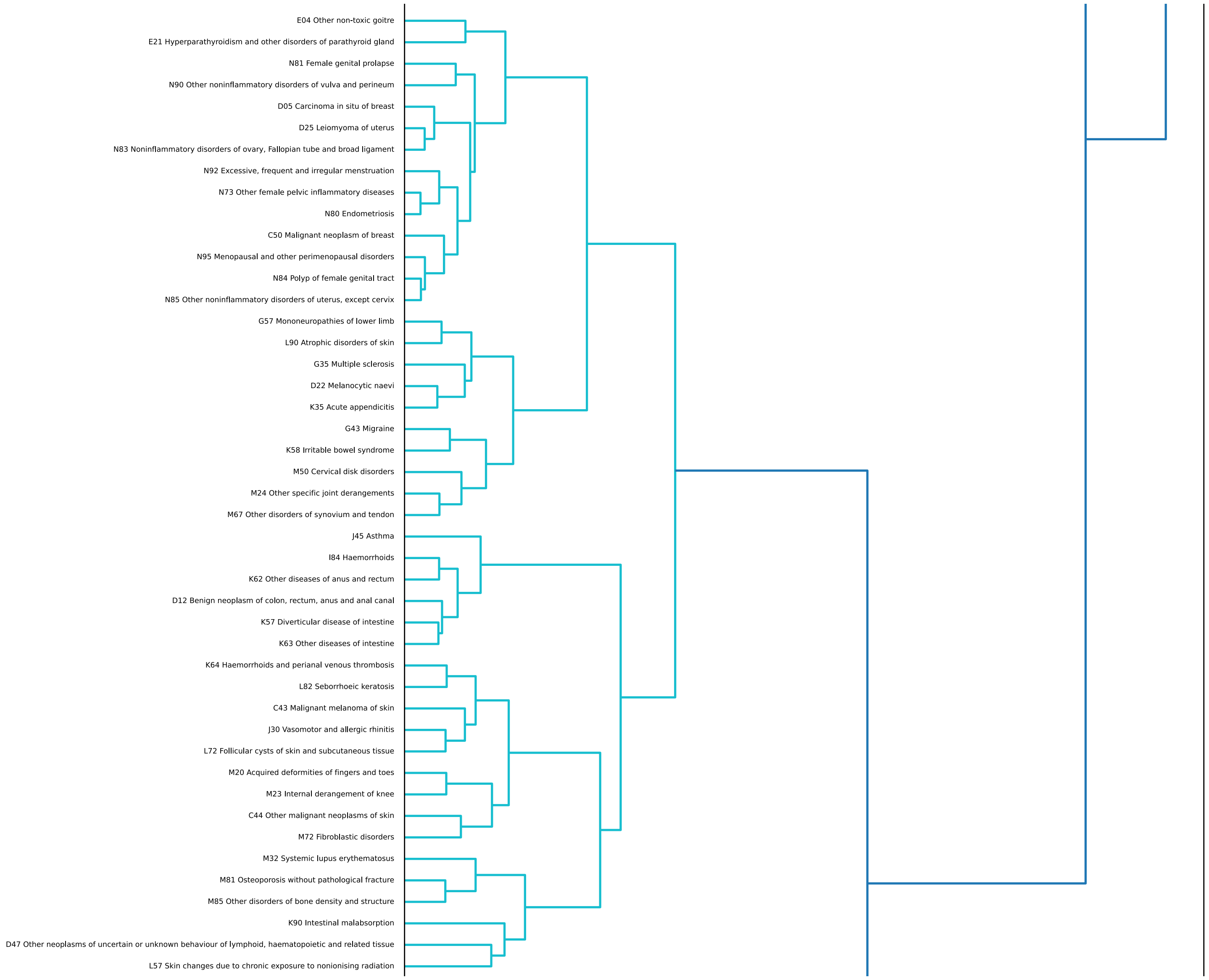
E04 Other non-toxic goitre
E21 Hyperparathyroidism and other disorders of parathyroid gland
N81 Female genital prolapse
N90 Other noninflammatory disorders of vulva and perineum
D05 Carcinoma in situ of breast
D25 Leiomyoma of uterus
N83 Noninflammatory disorders of ovary, Fallopian tube and broad ligament
N92 Excessive, frequent and irregular menstruation
N73 Other female pelvic inflammatory diseases
N80 Endometriosis
C50 Malignant neoplasm of breast
N95 Menopausal and other perimenopausal disorders
N84 Polyp of female genital tract
N85 Other noninflammatory disorders of uterus, except cervix
G57 Mononeuropathies of lower limb
L90 Atrophic disorders of skin
G35 Multiple sclerosis
D22 Melanocytic naevi
K35 Acute appendicitis
G43 Migraine
K58 Irritable bowel syndrome
M50 Cervical disk disorders
M24 Other specific joint derangements
M67 Other disorders of synovium and tendon
J45 Asthma
I84 Haemorrhoids
K62 Other diseases of anus and rectum
D12 Benign neoplasm of colon, rectum, anus and anal canal
K57 Diverticular disease of intestine
K63 Other diseases of intestine
K64 Haemorrhoids and perianal venous thrombosis
L82 Seborrhoeic keratosis
C43 Malignant melanoma of skin
J30 Vasomotor and allergic rhinitis
L72 Follicular cysts of skin and subcutaneous tissue
M20 Acquired deformities of fingers and toes
M23 Internal derangement of knee
C44 Other malignant neoplasms of skin
M72 Fibroblastic disorders
M32 Systemic lupus erythematosus
M81 Osteoporosis without pathological fracture
M85 Other disorders of bone density and structure
K90 Intestinal malabsorption
D47 Other neoplasms of uncertain or unknown behaviour of lymphoid, haematopoietic and related tissue
L57 Skin changes due to chronic exposure to nonionising radiation

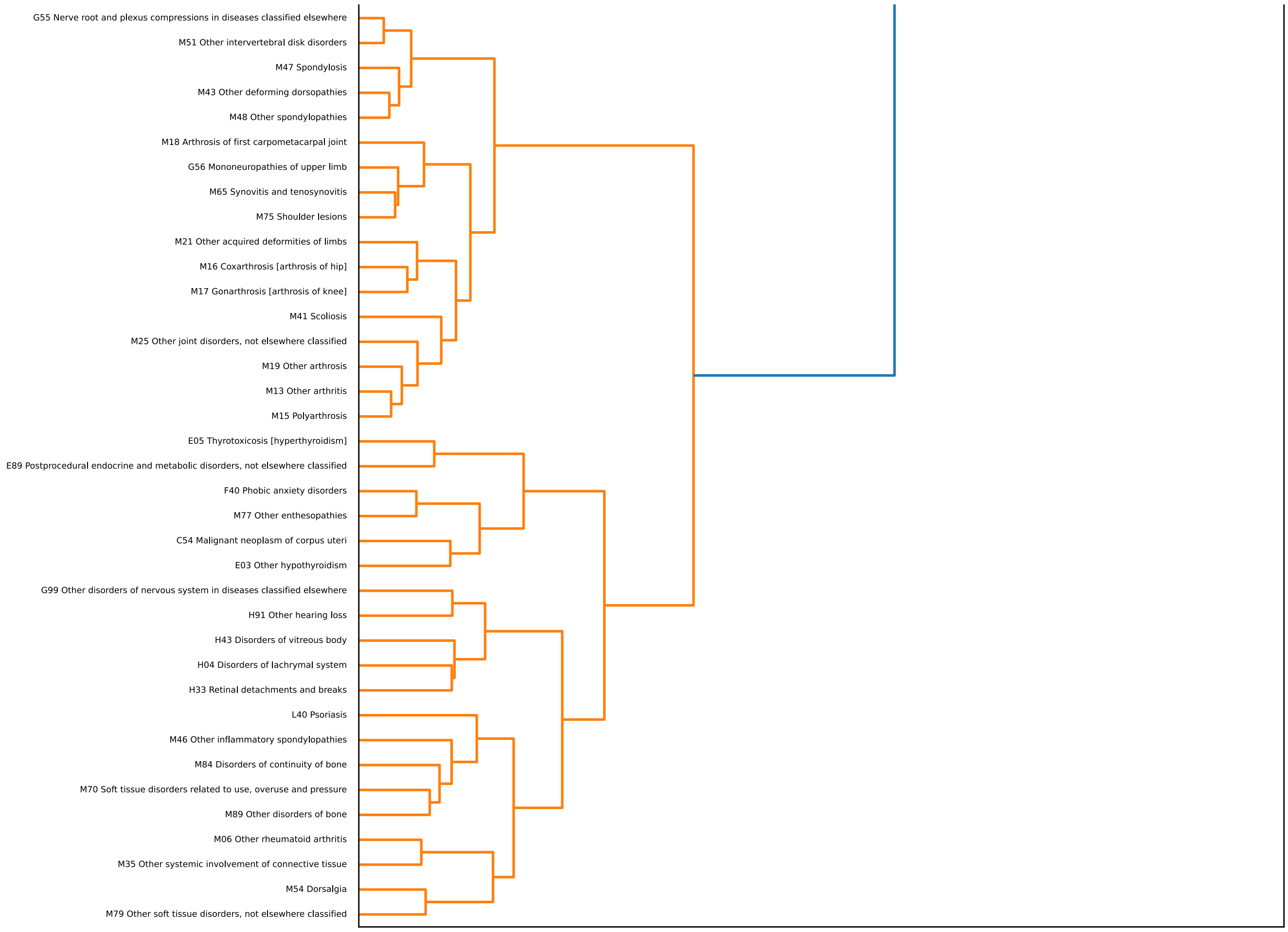


Figure 6: Dendrogram of clusters of $\mathbf{V}$.

#### Identifying the most important proteins in each cluster

To identify the most important proteins in each cluster, we first calculate the submatrix $\mathbf{B}_c$, $c = 1, \ldots, C$ for each of the $C$ clusters:

$$\mathbf{B}_c = \mathbf{V}_c \mathbf{W} \in \mathbb{R}^{D_c \times P}, \quad c = 1, \ldots, C,$$

where $\mathbf{V}_c$ is the submatrix of $\mathbf{V}$ containing only the diseases in cluster $c$ and $D_c$ is the number of diseases in cluster $c$. We then calculate the protein importance score $\tilde{s}_{cp}$ for each cluster as

$$\tilde{s}_{cp} = \frac{1}{D_c} \sum_{d=1}^{D_c} |b_{c_{dp}}|, \quad c = 1, \ldots, C, \; p = 1, \ldots, P.$$

The proteins in each cluster are then ranked by protein importance score.

### Clustering of $\mathbf{V}\mathbf{V}^\top$

The matrix $\mathbf{V}\mathbf{V}^\top$ was clustered using Ward's method for agglomerative clustering and the number is clusters was set to 7 based on the elbow method. The corresponding elbow plot can be seen in Figure 7.

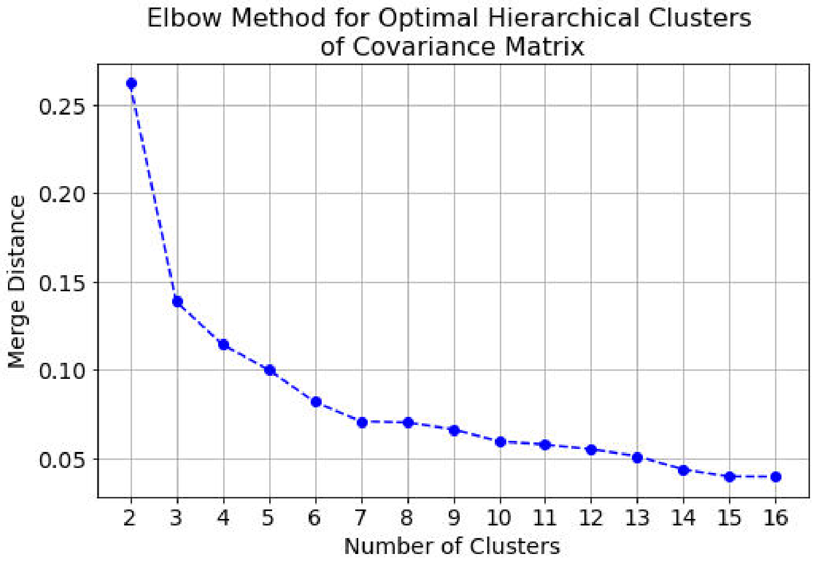


Figure 7: Elbow plot for the clusters of $\mathbf{V}\mathbf{V}^\top$.

The full dendrogram with disease labels from Figure 4 is shown in Figure 8:

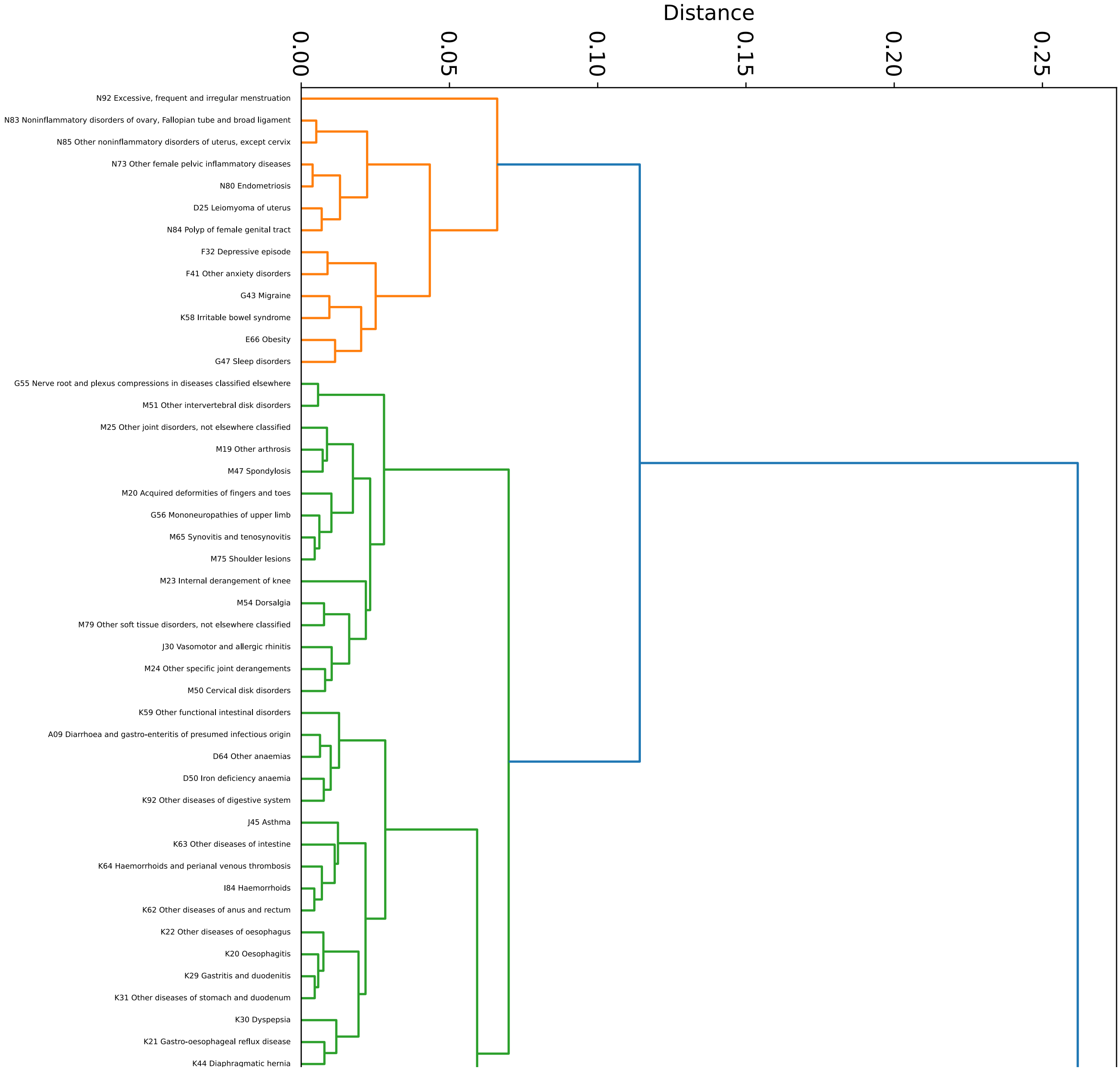

Distance
0.00
0.05
0.10
0.15
0.20
0.25
N92 Excessive, frequent and irregular menstruation
N83 Noninflammatory disorders of ovary, Fallopian tube and broad ligament
N85 Other noninflammatory disorders of uterus, except cervix
N73 Other female pelvic inflammatory diseases
N80 Endometriosis
D25 Leiomyoma of uterus
N84 Polyp of female genital tract
F32 Depressive episode
F41 Other anxiety disorders
G43 Migraine
K58 Irritable bowel syndrome
E66 Obesity
G47 Sleep disorders
G55 Nerve root and plexus compressions in diseases classified elsewhere
M51 Other intervertebral disk disorders
M25 Other joint disorders, not elsewhere classified
M19 Other arthrosis
M47 Spondylosis
M20 Acquired deformities of fingers and toes
G56 Mononeuropathies of upper limb
M65 Synovitis and tenosynovitis
M75 Shoulder lesions
M23 Internal derangement of knee
M54 Dorsalgia
M79 Other soft tissue disorders, not elsewhere classified
J30 Vasomotor and allergic rhinitis
M24 Other specific joint derangements
M50 Cervical disk disorders
K59 Other functional intestinal disorders
A09 Diarrhoea and gastro-enteritis of presumed infectious origin
D64 Other anaemias
D50 Iron deficiency anaemia
K92 Other diseases of digestive system
J45 Asthma
K63 Other diseases of intestine
K64 Haemorrhoids and perianal venous thrombosis
I84 Haemorrhoids
K62 Other diseases of anus and rectum
K22 Other diseases of oesophagus
K20 Oesophagitis
K29 Gastritis and duodenitis
K31 Other diseases of stomach and duodenum
K30 Dyspepsia
K21 Gastro-oesophageal reflux disease
K44 Diaphragmatic hernia

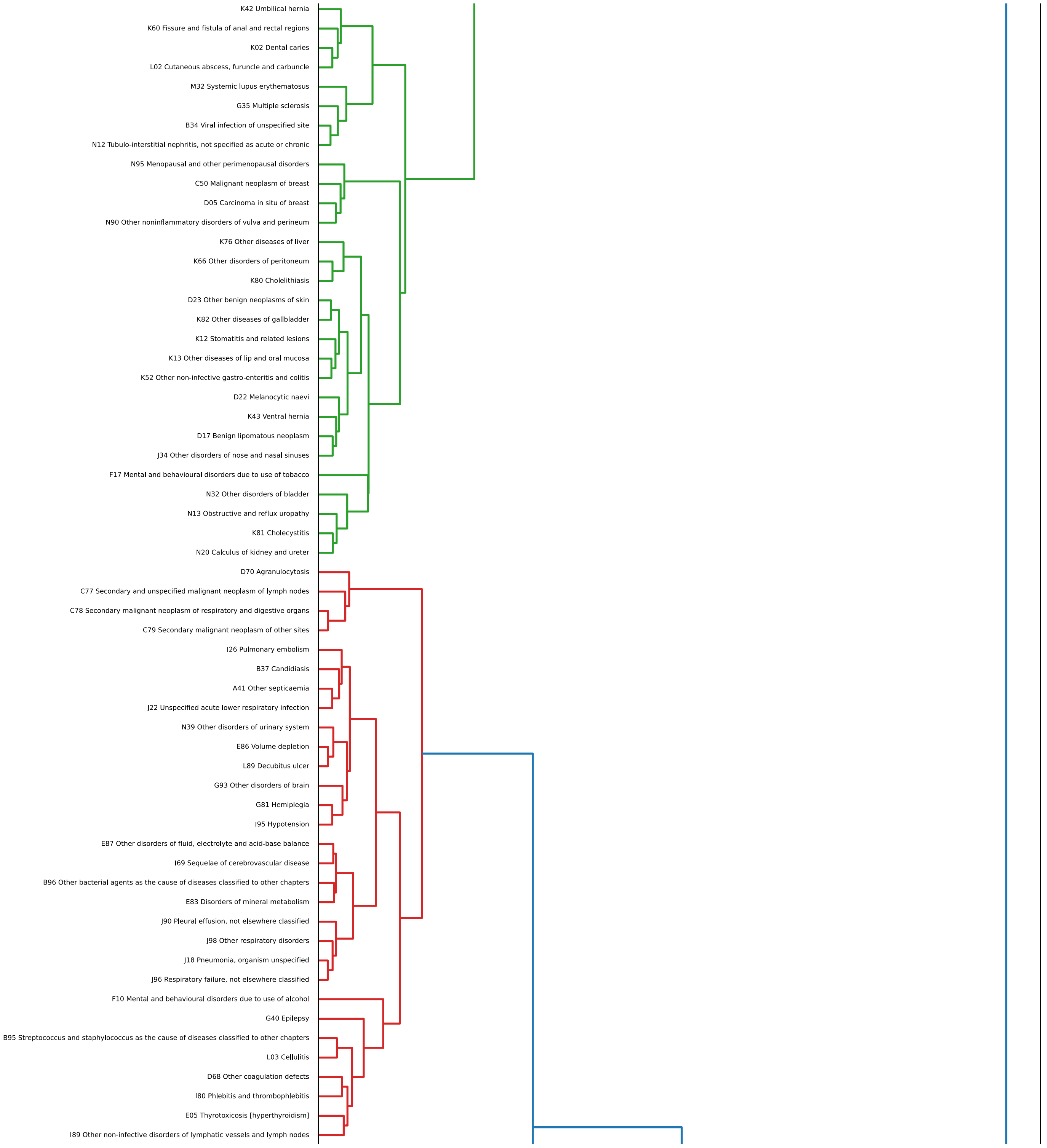

K42 Umbilical hernia
K60 Fissure and fistula of anal and rectal regions
K02 Dental caries
L02 Cutaneous abscess, furuncle and carbuncle
M32 Systemic lupus erythematosus
G35 Multiple sclerosis
B34 Viral infection of unspecified site
N12 Tubulo-interstitial nephritis, not specified as acute or chronic
N95 Menopausal and other perimenopausal disorders
C50 Malignant neoplasm of breast
D05 Carcinoma in situ of breast
N90 Other noninflammatory disorders of vulva and perineum
K76 Other diseases of liver
K66 Other disorders of peritoneum
K80 Cholelithiasis
D23 Other benign neoplasms of skin
K82 Other diseases of gallbladder
K12 Stomatitis and related lesions
K13 Other diseases of lip and oral mucosa
K52 Other non-infective gastro-enteritis and colitis
D22 Melanocytic naevi
K43 Ventral hernia
D17 Benign lipomatous neoplasm
J34 Other disorders of nose and nasal sinuses
F17 Mental and behavioural disorders due to use of tobacco
N32 Other disorders of bladder
N13 Obstructive and reflux uropathy
K81 Cholecystitis
N20 Calculus of kidney and ureter
D70 Agranulocytosis
C77 Secondary and unspecified malignant neoplasm of lymph nodes
C78 Secondary malignant neoplasm of respiratory and digestive organs
C79 Secondary malignant neoplasm of other sites
I26 Pulmonary embolism
B37 Candidiasis
A41 Other septicaemia
J22 Unspecified acute lower respiratory infection
N39 Other disorders of urinary system
E86 Volume depletion
L89 Decubitus ulcer
G93 Other disorders of brain
G81 Hemiplegia
I95 Hypotension
E87 Other disorders of fluid, electrolyte and acid-base balance
I69 Sequelae of cerebrovascular disease
B96 Other bacterial agents as the cause of diseases classified to other chapters
E83 Disorders of mineral metabolism
J90 Pleural effusion, not elsewhere classified
J98 Other respiratory disorders
J18 Pneumonia, organism unspecified
J96 Respiratory failure, not elsewhere classified
F10 Mental and behavioural disorders due to use of alcohol
G40 Epilepsy
B95 Streptococcus and staphylococcus as the cause of diseases classified to other chapters
L03 Cellulitis
D68 Other coagulation defects
I80 Phlebitis and thrombophlebitis
E05 Thyrotoxicosis [hyperthyroidism]
I89 Other non-infective disorders of lymphatic vessels and lymph nodes

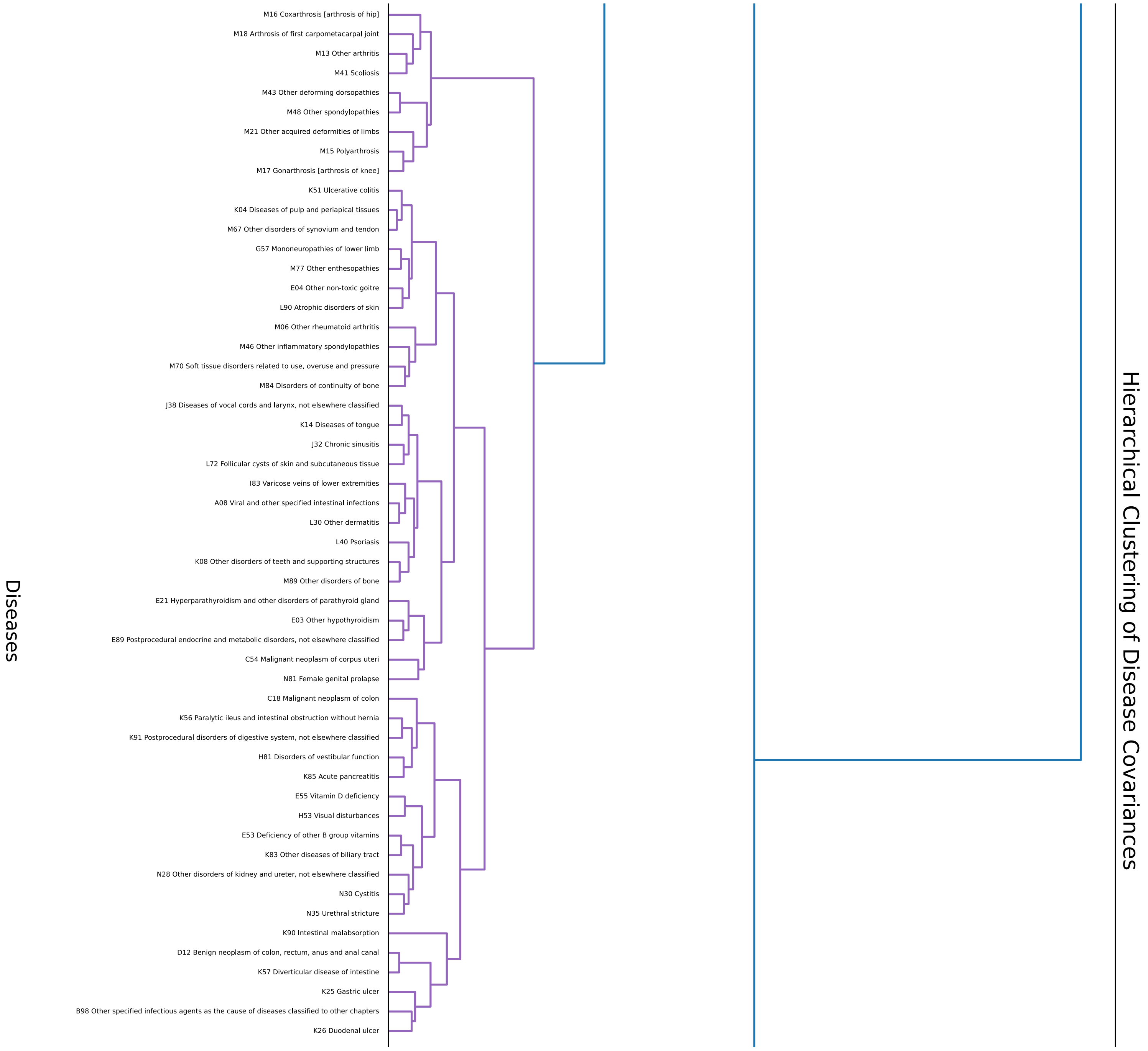
Hierarchical Clustering of Disease Covariances
Diseases
M16 Coxarthrosis [arthrosis of hip]
M18 Arthrosis of first carpometacarpal joint
M13 Other arthritis
M41 Scoliosis
M43 Other deforming dorsopathies
M48 Other spondylopathies
M21 Other acquired deformities of limbs
M15 Polyarthrosis
M17 Gonarthrosis [arthrosis of knee]
K51 Ulcerative colitis
K04 Diseases of pulp and periapical tissues
M67 Other disorders of synovium and tendon
G57 Mononeuropathies of lower limb
M77 Other enthesopathies
E04 Other non-toxic goitre
L90 Atrophic disorders of skin
M06 Other rheumatoid arthritis
M46 Other inflammatory spondylopathies
M70 Soft tissue disorders related to use, overuse and pressure
M84 Disorders of continuity of bone
J38 Diseases of vocal cords and larynx, not elsewhere classified
K14 Diseases of tongue
J32 Chronic sinusitis
L72 Follicular cysts of skin and subcutaneous tissue
I83 Varicose veins of lower extremities
A08 Viral and other specified intestinal infections
L30 Other dermatitis
L40 Psoriasis
K08 Other disorders of teeth and supporting structures
M89 Other disorders of bone
E21 Hyperparathyroidism and other disorders of parathyroid gland
E03 Other hypothyroidism
E89 Postprocedural endocrine and metabolic disorders, not elsewhere classified
C54 Malignant neoplasm of corpus uteri
N81 Female genital prolapse
C18 Malignant neoplasm of colon
K56 Paralytic ileus and intestinal obstruction without hernia
K91 Postprocedural disorders of digestive system, not elsewhere classified
H81 Disorders of vestibular function
K85 Acute pancreatitis
E55 Vitamin D deficiency
H53 Visual disturbances
E53 Deficiency of other B group vitamins
K83 Other diseases of biliary tract
N28 Other disorders of kidney and ureter, not elsewhere classified
N30 Cystitis
N35 Urethral stricture
K90 Intestinal malabsorption
D12 Benign neoplasm of colon, rectum, anus and anal canal
K57 Diverticular disease of intestine
K25 Gastric ulcer
B98 Other specified infectious agents as the cause of diseases classified to other chapters
K26 Duodenal ulcer

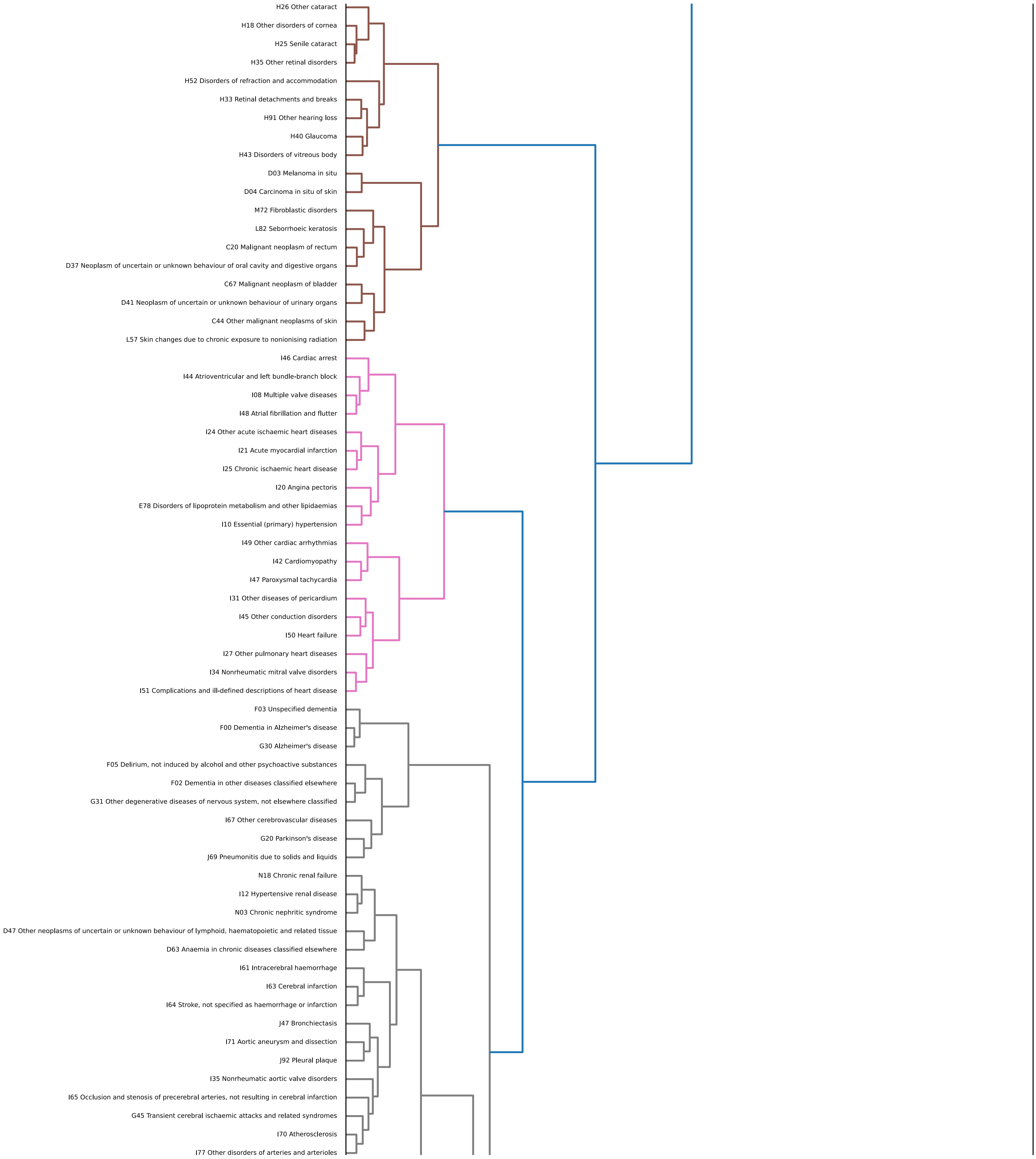
H26 Other cataract
H18 Other disorders of cornea
H25 Senile cataract
H35 Other retinal disorders
H52 Disorders of refraction and accommodation
H33 Retinal detachments and breaks
H91 Other hearing loss
H40 Glaucoma
H43 Disorders of vitreous body
D03 Melanoma in situ
D04 Carcinoma in situ of skin
M72 Fibroblastic disorders
L82 Seborrhoeic keratosis
C20 Malignant neoplasm of rectum
D37 Neoplasm of uncertain or unknown behaviour of oral cavity and digestive organs
C67 Malignant neoplasm of bladder
D41 Neoplasm of uncertain or unknown behaviour of urinary organs
C44 Other malignant neoplasms of skin
L57 Skin changes due to chronic exposure to nonionising radiation
I46 Cardiac arrest
I44 Atrioventricular and left bundle-branch block
I08 Multiple valve diseases
I48 Atrial fibrillation and flutter
I24 Other acute ischaemic heart diseases
I21 Acute myocardial infarction
I25 Chronic ischaemic heart disease
I20 Angina pectoris
E78 Disorders of lipoprotein metabolism and other lipidaemias
I10 Essential (primary) hypertension
I49 Other cardiac arrhythmias
I42 Cardiomyopathy
I47 Paroxysmal tachycardia
I31 Other diseases of pericardium
I45 Other conduction disorders
I50 Heart failure
I27 Other pulmonary heart diseases
I34 Nonrheumatic mitral valve disorders
I51 Complications and ill-defined descriptions of heart disease
F03 Unspecified dementia
F00 Dementia in Alzheimer's disease
G30 Alzheimer's disease
F05 Delirium, not induced by alcohol and other psychoactive substances
F02 Dementia in other diseases classified elsewhere
G31 Other degenerative diseases of nervous system, not elsewhere classified
I67 Other cerebrovascular diseases
G20 Parkinson's disease
J69 Pneumonitis due to solids and liquids
N18 Chronic renal failure
I12 Hypertensive renal disease
N03 Chronic nephritic syndrome
D47 Other neoplasms of uncertain or unknown behaviour of lymphoid, haematopoietic and related tissue
D63 Anaemia in chronic diseases classified elsewhere
I61 Intracerebral haemorrhage
I63 Cerebral infarction
I64 Stroke, not specified as haemorrhage or infarction
J47 Bronchiectasis
I71 Aortic aneurysm and dissection
J92 Pleural plaque
I35 Nonrheumatic aortic valve disorders
I65 Occlusion and stenosis of precerebral arteries, not resulting in cerebral infarction
G45 Transient cerebral ischaemic attacks and related syndromes
I70 Atherosclerosis
I77 Other disorders of arteries and arterioles

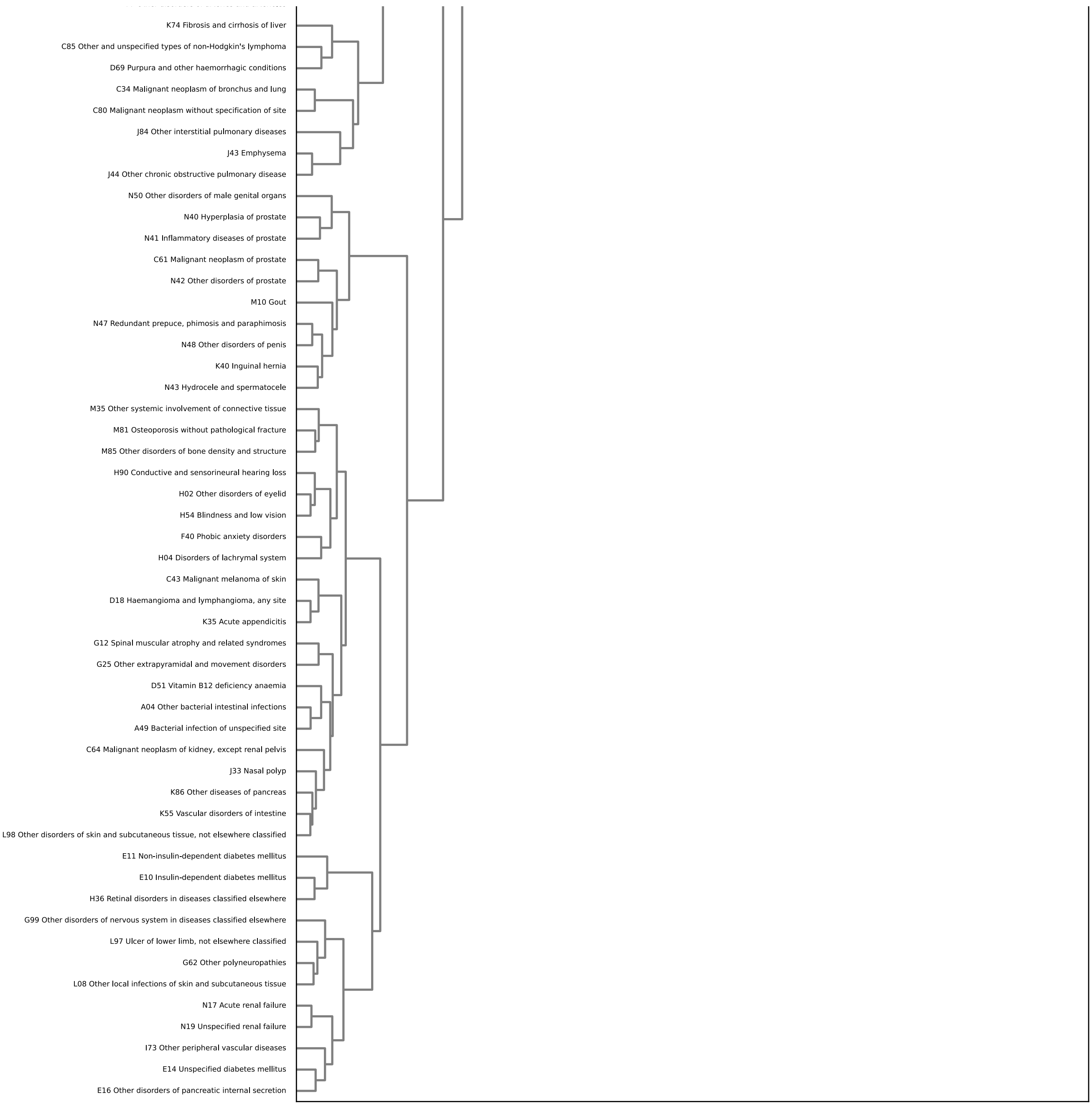


Figure 8: Dendrogram of clusters of $\mathbf{V}\mathbf{V}^\top$.

### Influence of the proteins in the minimal panel on diseases

Figure 9 shows the influence of the 10 proteins included in the panel on the diseases included in the analysis.

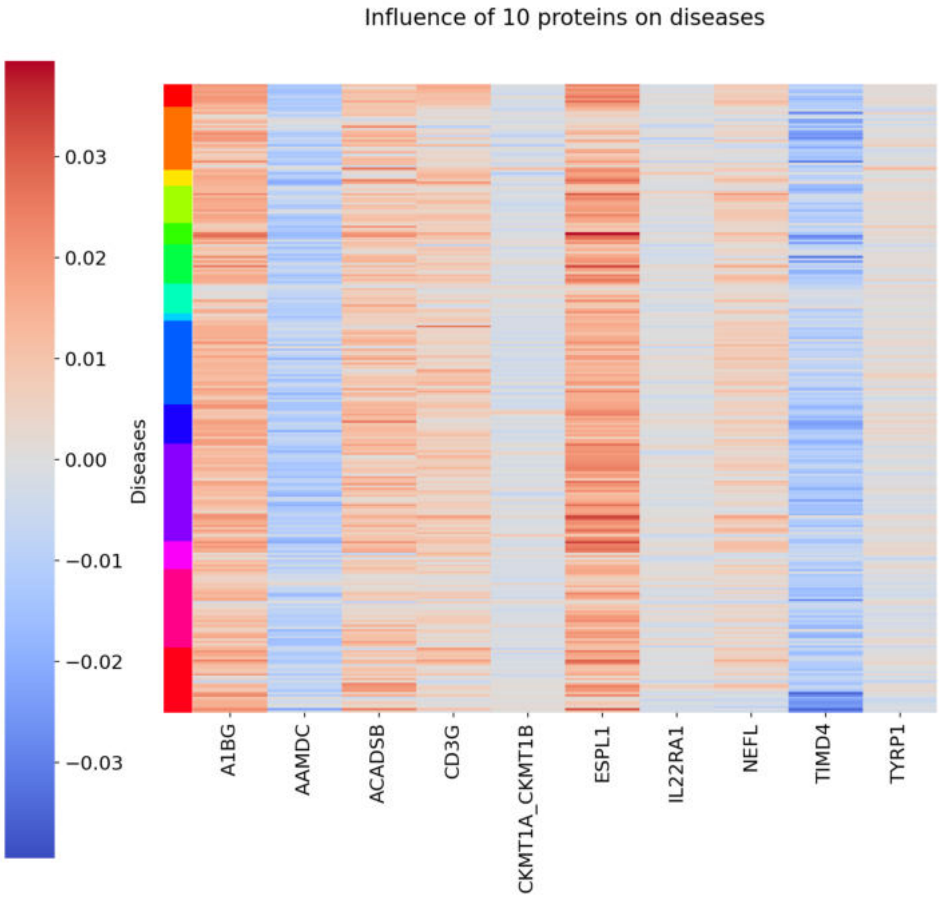


Figure 9: Influence of each of the 10 proteins in the minimal panel on disease risk.

### Kaplan-Meier trajectories across different risk groups

We want to analyze the disease risk in a different way by plotting the empirical probability of getting a disease by calculating 1- the Kaplan-Meier estimator (Kaplan and Meier, 1958) for different risk groups, similar to the analyses of cumulative incidence plots in (Argentieri et al., 2024). For each disease, the high risk group is defined as the decile of UK Biobank participants with the highest rate of ageing, the medium risk group consists of the patients whose rates of ageing are in the middle decile and the low risk group contains the patients with the $10\%$ lowest rates of ageing. These plots are shown in figure 10.

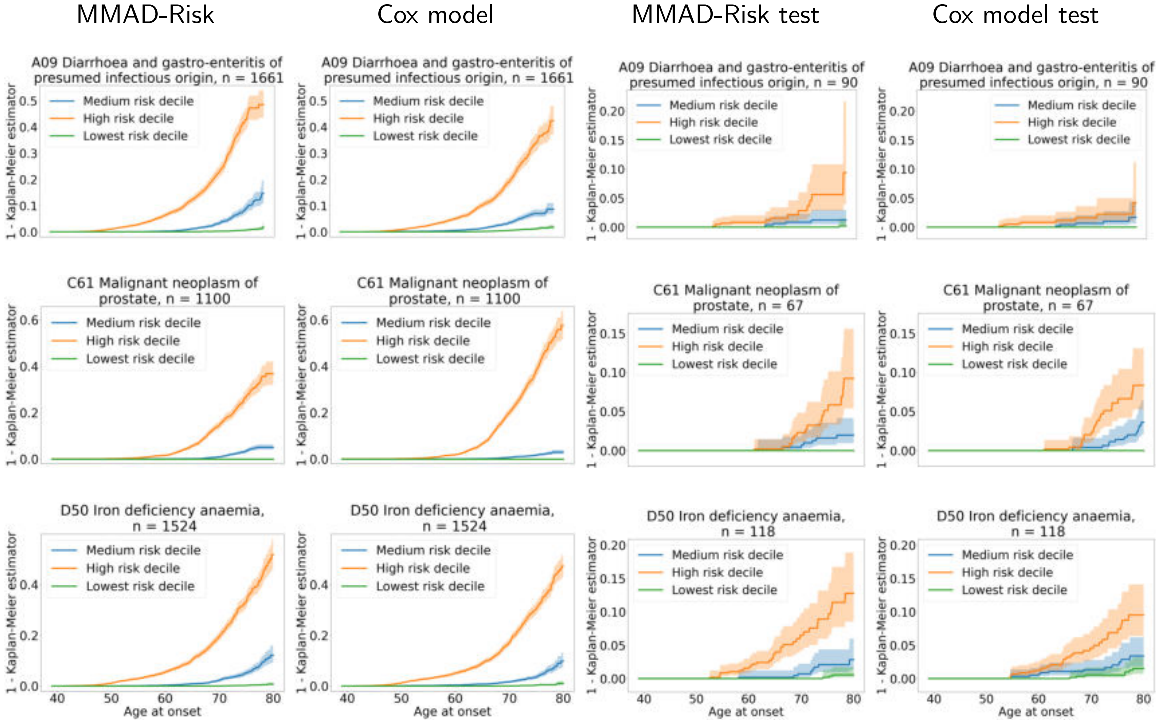


Figure 10: Stratified plots of 1-Kaplan Meier estimator for MMAD-Risk and the Cox proportional hazards model. The left column shows the stratified survival curves for MMAD-Risk on the entire data set and the second column on the left the results for the Cox proportional hazards model on the entire data set. The third column from left shows the results for MMAD-Risk on the test data set, the last column the results for the Cox proportional hazards model on the test data set. The orange curve represents the participants in the highest risk decile, the blue curve represents the $10\%$ participants with median risk and the green represents the participants in the lowest risk decile.

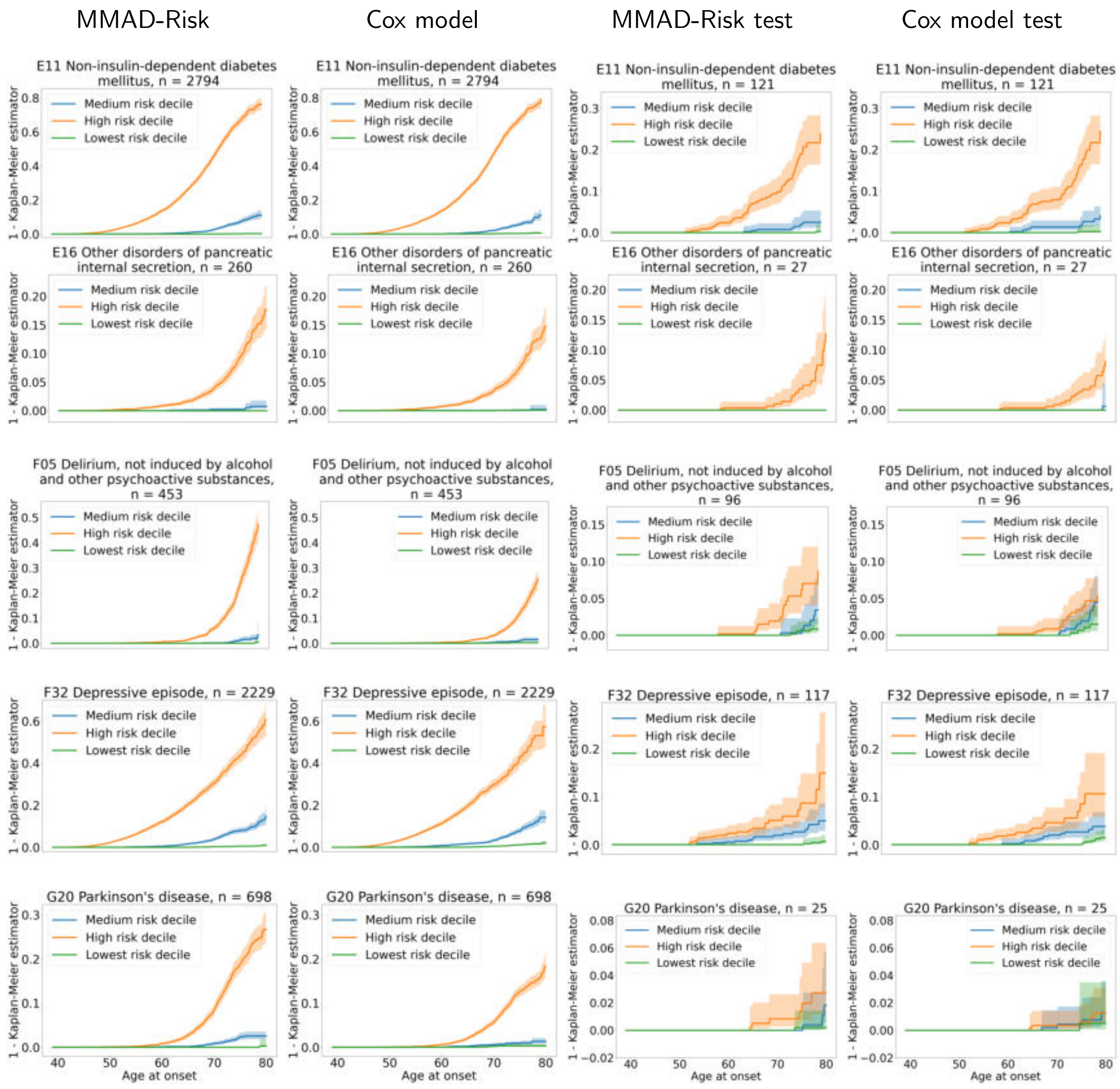


Figure 10: Stratified plots of 1-Kaplan Meier estimator for MMAD-Risk and the Cox proportional hazards model. The left column shows the stratified survival curves for MMAD-Risk on the entire data set and the second column on the left the results for the Cox proportional hazards model on the entire data set. The third column from left shows the results for MMAD-Risk on the test data set, the last column the results for the Cox proportional hazards model on the test data set. The orange curve represents the participants in the highest risk decile, the blue curve represents the $10\%$ participants with median risk and the green represents the participants in the lowest risk decile.

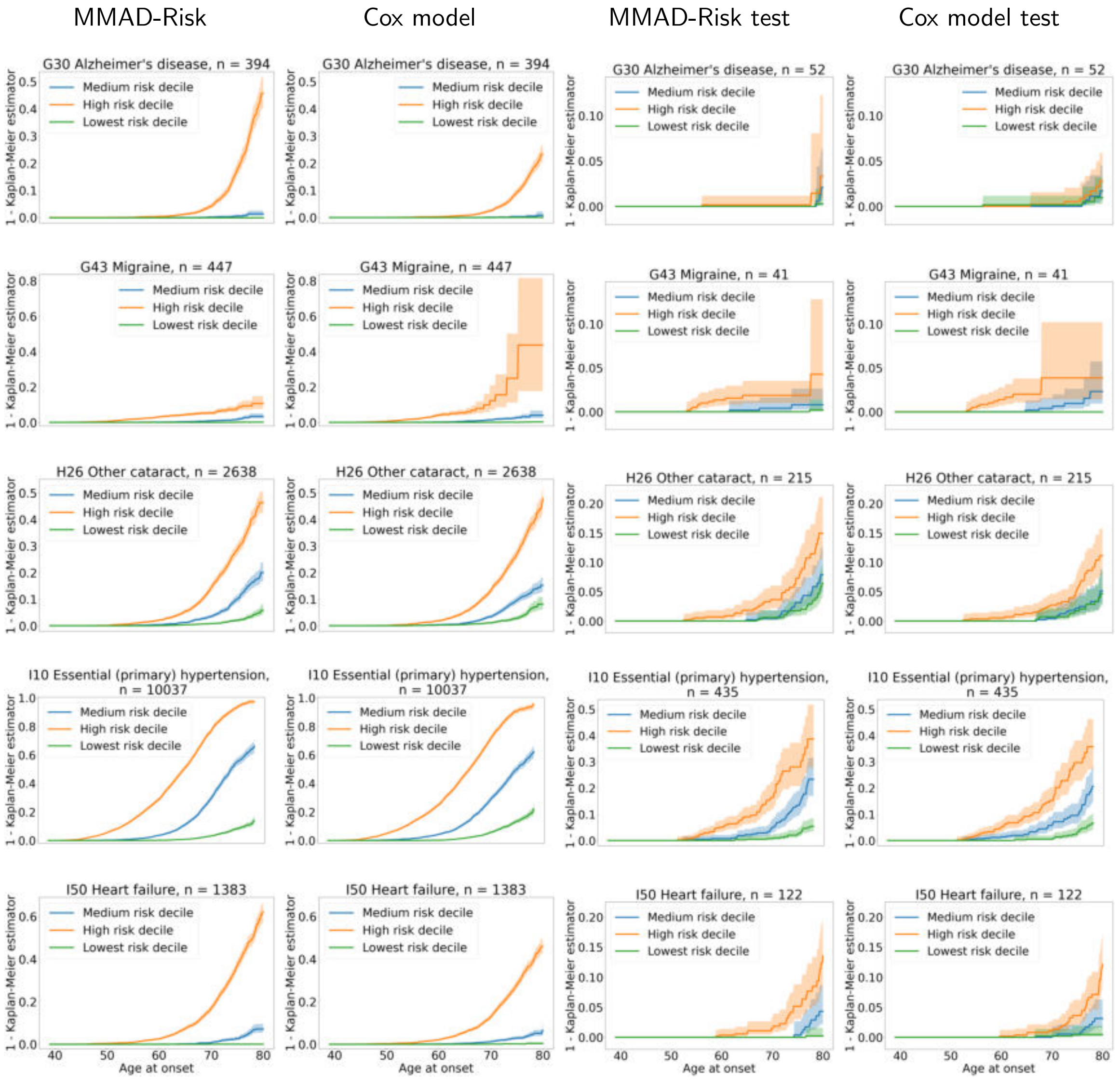


Figure 10: Stratified plots of 1-Kaplan Meier estimator for MMAD-Risk and the Cox proportional hazards model. The left column shows the stratified survival curves for MMAD-Risk on the entire data set and the second column on the left the results for the Cox proportional hazards model on the entire data set. The third column from left shows the results for MMAD-Risk on the test data set, the last column the results for the Cox proportional hazards model on the test data set. The orange curve represents the participants in the highest risk decile, the blue curve represents the $10\%$ participants with median risk and the green represents the participants in the lowest risk decile.

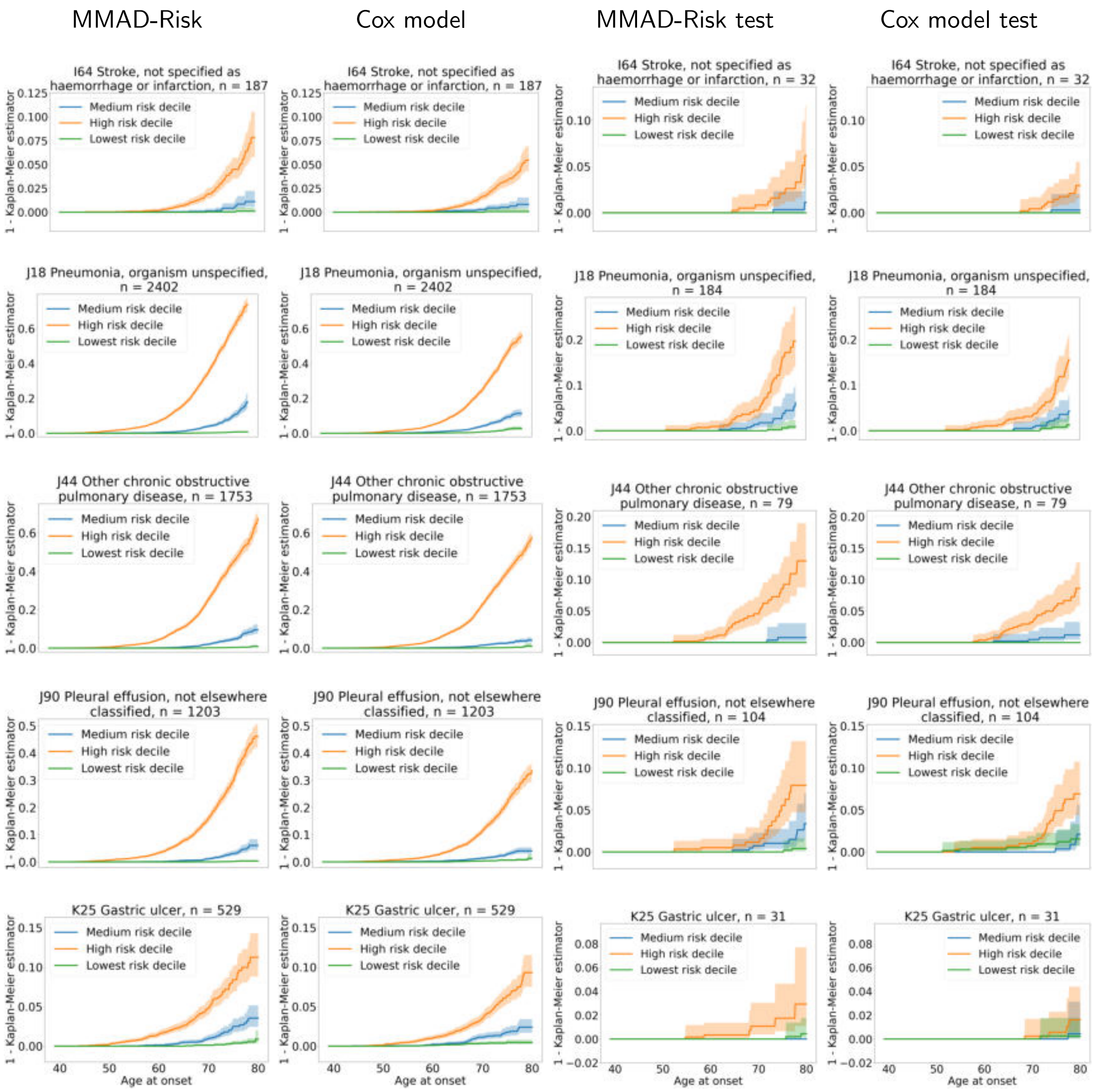


Figure 10: Stratified plots of 1-Kaplan Meier estimator for MMAD-Risk and the Cox proportional hazards model. The left column shows the stratified survival curves for MMAD-Risk on the entire data set and the second column on the left the results for the Cox proportional hazards model on the entire data set. The third column from left shows the results for MMAD-Risk on the test data set, the last column the results for the Cox proportional hazards model on the test data set. The orange curve represents the participants in the highest risk decile, the blue curve represents the $10\%$ participants with median risk and the green represents the participants in the lowest risk decile.

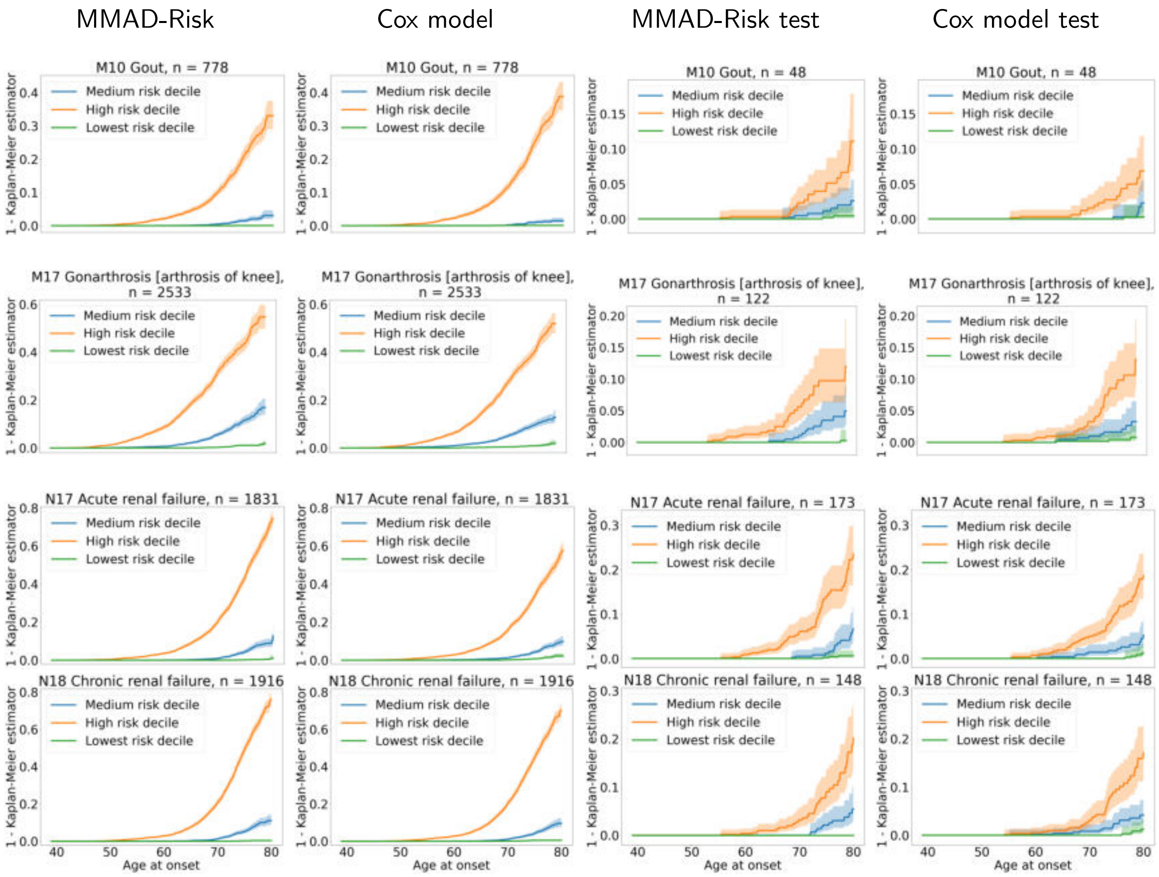


Figure 10: Stratified plots of 1-Kaplan Meier estimator for MMAD-Risk and the Cox proportional hazards model. The left column shows the stratified survival curves for MMAD-Risk on the entire data set and the second column on the left the results for the Cox proportional hazards model on the entire data set. The third column from left shows the results for MMAD-Risk on the test data set, the last column the results for the Cox proportional hazards model on the test data set. The orange curve represents the participants in the highest risk decile, the blue curve represents the $10\%$ participants with median risk and the green represents the participants in the lowest risk decile.

### Declaration on the use of Large Language Models (LLMs)

During the preparation of this work, AMH used Qwen-3-30B-A3B-Instruct and Claude Sonnet 4.6 and 4.7 to assist in writing and debugging the software. AMH reviewed, tested, and optimized all generated code independently and take full responsibility for the accuracy of the computational results.

AMH used Qwen-3-30B-A3B-Instruct 2507 proofreading the manuscript and to suggest rephrasing for selected sentences in the abstract, the methods section and the discussion. All content was critically read, revised, and validated by AMH and JS, who maintain full accountability for the scientific integrity of the publication.
